# Temporal Control of CRISPR-Cas9 Activity Using Bio-orthogonal Chemistry

**DOI:** 10.1101/2025.07.01.662593

**Authors:** Bhoomika Pandit, Sweta Vangaveti, Justa Fidelity Sentre, Gabriele Fuchs, Maksim Royzen

## Abstract

The CRISPR–Cas9 system has become a widely used tool for genome engineering. Here we present a new method for small-molecule control of CRISPR-Cas9 using bio-orthogonal chemistry between tetrazine (Tz) and *trans*-cyclooctene (TCO). We carried out molecular modeling studies and identified a unique position on single guide RNA (sgRNA) that can be site-specifically tagged with Tz without disrupting its activity. We also synthesized a series of TCO-modified CRISPR suppressors. When exogenously added, they click to the Tz-tagged sgRNA, perturb the system and drastically reduce the nuclease activity. The most successful suppressor is a TCO-modified six amino acid long cell-penetrating peptide, which shows excellent cell permeability. We showed that out method to control CRISPR-Cas9 nuclease activity is general by applying it to three different sgRNAs. We also showed that our method works in solution, as well as live HEL293 cells. We utilized flow cytometry to demonstrate temporal control of CRISPR-Cas9 targeting GFP. Lastly, we showed the therapeutic potential of our method by targeting vascular endothelial growth factor A (VEGFA).

## Introduction

The CRISPR (clustered regularly interspaced short palindromic repeats)–Cas9 system has become a widely popular tool for genome engineering in different organisms and biological systems. The most frequently used system is the type II Cas9 from *Streptococcus pyogenes* strain SF370 (SpyCas9) and single guide RNA (sgRNA), which targets specific DNA sequences in the genome to create a blunt-ended double-strand breaks.^1,2^ In recent years, CRISPR technology has been evaluated in several human clinical trials to treat various cancers, eye disease and chronic infection.^3–5^ Perhaps the most striking biomedical breakthrough has been in the treatment of sickle cell disease, the most common inherited hemoglobinopathy worldwide.^6,7^ In 2023, FDA approve two milestone sickle cell disease treatments, Casgevy and Lyfgenia, that are based on CRISPR technology.

One of the biggest hurdles in the development of new CRISPR-Cas9-based drugs is elimination of the so-called off-target effects. After administration, sgRNA-Cas9 ribonuclear protein will stay in the body for a long time. In addition to the intended target gene, CRISPR-Cas9-induced indels can occur at unintended off-target cleavage sites that have as many as five mismatches within the protospacer.^8^ Editing of off-target sites is known to occur on a slower timescale than the on-target sites.^7,9,10^ Therefore, there is considerable interest in developing systems that allow temporal control CRISPR-Cas9 nuclease activity. Such systems will inactivate either Cas9 or sgRNA once the target gene has been edited to prevent further editing of off-target sites.

A number of bio-engineering approaches have already been reported to modulate activity of Cas9. For example, Choudhary group fused Cas9 to the destabilized polypeptide domains of *E. coli’s* dihydrofolate reductase.^11^ These domains are unstable and rapidly target the fusion protein for proteasomal degradation. The destabilized domains can be stabilized upon addition of trimethoprim compound, which prevents proteasomal degradation. Choudhary group have shown that by controlling the concentration and time of administration of trimethoprim, both the dosage and temporal control of Cas9 activity can be achieved. Alternative methods to modulate Cas9 activity have been reported by Bondy-Denomy and Yee groups.^12–14^

Alternatively, CRISPR-Cas9 nuclease activity can be controlled by engineering ligand-responsive sgRNA. Batey and co-workers fused theophylline-binding aptamer into the sgRNA’s tetraloop, which is known to be highly tolerant of nucleotide insertions.^15^ The authors subsequently introduced fourteen nucleotide mutations and carried out a selection process that yielded constructs that exhibited nuclease activity only in the presence of theophylline, which binds the aptamer with nanomolar affinity. Kundert *et al.* reported a similar system containing sgRNA-fused aptamer which can be controlled by small molecule ligands.^16^ An alternative method to control sgRNA activity has been described by Tian.^17^ All of the described systems are synthetically complex and require considerable bio-engineering with major perturbation of the native CRISPR-Cas9 system.

Herein we describe an approach to control CRISPR-Cas9 activity using small molecule bio-orthogonal chemistry between *trans*-cyclooctene (TCO) and tetrazine (Tz). The reaction between TCO and Tz has exceptionally fast kinetics. The two reacting groups are biocompatible and have high potential for bio-medical translation.^18,19^ The design is schematically illustrated in **Figure 1**. In our design sgRNA is modified with a covalently attached Tz tag. Tz tagging of a single nucleotide has been experimentally optimized to cause minimal perturbation to sgRNA’s function in Cas9-assisted nuclease activity. Thus, the construct shown on the left in **Figure 1** is constitutively functional. To control CRISPR activity, TCO-containing small molecule suppressor will be exogenously added. The TCO group of the suppressor will react via bio-orthogonal click reaction with the Tz group attached to sgRNA. The conjugation product will cause perturbation in the domain of sgRNA/Cas9 complex that is important for nuclease activity, thus rendering the RNP complex inactive. In comparison to already reported methods to control CRISPR-Cas9 activity, the key advantage of our design is that it involves a small molecule RNA tag, which minimally perturbs sgRNA’s structure, and small molecule suppressors which can be easily administered to control the nuclease activity.

**Figure 1.**
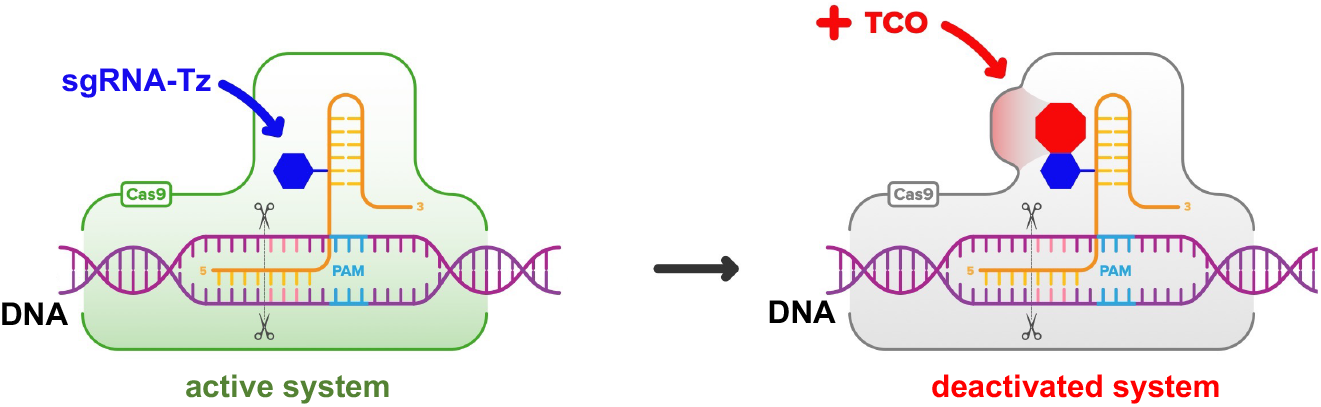
Schematic representation of controlled turn-off of CRISPR-Cas9 activity using bio-orthogonal chemistry between TCO and Tz.

## Results and Discussion

### (A) Design of constitutively active Tz-tagged sgRNA

We decided to explore the repeat:anti-repeat region of sgRNA for covalent tagging with Tz (**Figure 2A**). Several studies showed that it is a highly sensitive and important area of the sgRNA-Cas9 ribonucleoprotein complex. Nishimasu *et al.* reported the crystal structure of Cas9 in complex with sgRNA and target DNA.^20^ Cas9 consists of two lobes: a recognition (REC) lobe and a nuclease (NUC) lobe. REC1 and REC2 domains of the REC lobe make direct contacts with the repeat:anti-repeat region of sgRNA. Deletion of either the repeat-interacting region or the anti-repeat-interacting region of the REC1 domain abolished the nuclease activity. Sontheimer *et al.* reported that 2′-OMe RNA modification of all uridines in the repeat:anti-repeat region significantly lowered nuclease activity.^21,22^ Langer and Anderson showed that replacement of the entire repeat:anti-repeat with the corresponding DNA nucleotides completely eliminated nuclease activity.^22^ Lastly, Gagnon and Damha reported that replacement of the entire repeat:anti-repeat region with 2′-F-ANA lead to an analogous outcome.^23^

**Figure 2.**
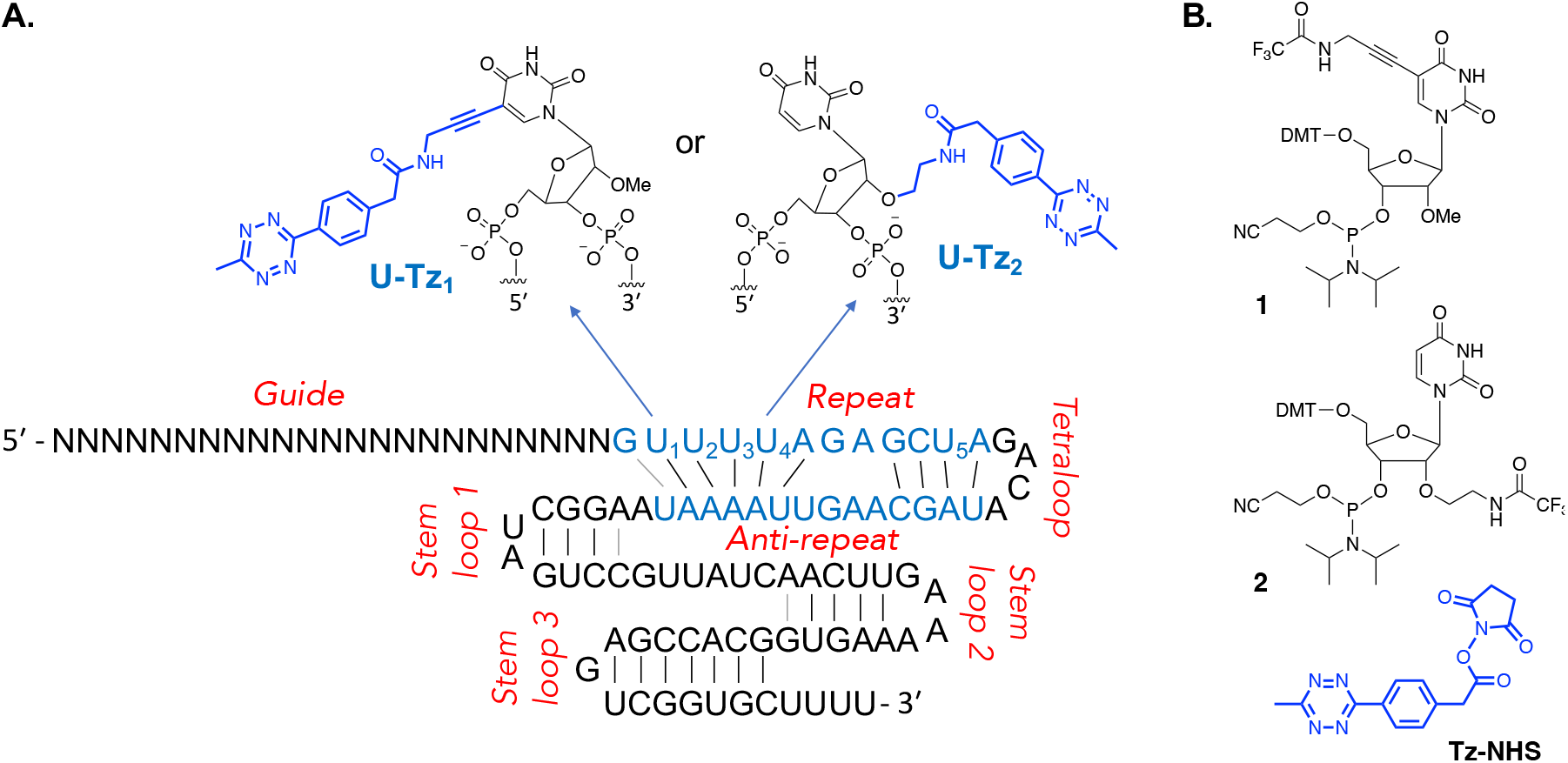
(**A.**) Incorporation of Tz into the repeat region of sgRNA. The important uridine residues are numbered U_1_-U_4_. (**B**) Phosphoramidites and Tz-NHS ester that were used to tag sgRNAs with Tz.

To understand the structural context and evaluate the feasibility of tagging the uridines in the repeat sequence of sgRNA with Tz, we carried out molecular modeling studies utilizing the experimentally solved crystal structure of Cas9:RNA/DNA complex.^20^ First, the solvent accessible surface area (SASA) of the uridines in the repeat sequence was calculated. This revealed that U_2_ and U_3_ have lower SASA compared to U_1_ and U_4_ (**Figure 3a**). U_4_ however, is closer to the peripheral surface of the enzyme compared to U_1_ (**Figure 3c**). Furthermore, the number of contacts with Cas9, defined as any atoms of the Us within 4Å of the enzyme, is lower for U_1_ and U_4_ compared to U_2_ and U_3_ (**Figure 3b**). While C5, the site for the Tz_1_ modification, has no contacts with the enzyme, the number of contacts for 2′-O, the site for the Tz_2_ modification, is higher in case of U_1_. Additionally modeling the Tz_1_ and Tz_2_ modifications on both U_1_ and U_4_ revealed that the modifications can be tolerated with minor local rearrangements and minimal disruption of the RNA:enzyme interface in the REC1 domain (**Figure 3d & 3e**). This is particularly true for U_4_, where the modified groups point away from the complex. Thus, the modeling predicted that while sgRNA modified with Tz_1_ or Tz_2_ at U_1_ and U_4_ should remain active, the modifications might be more accessible for accommodating additional chemistries at position U_4_.

**Figure 3.**
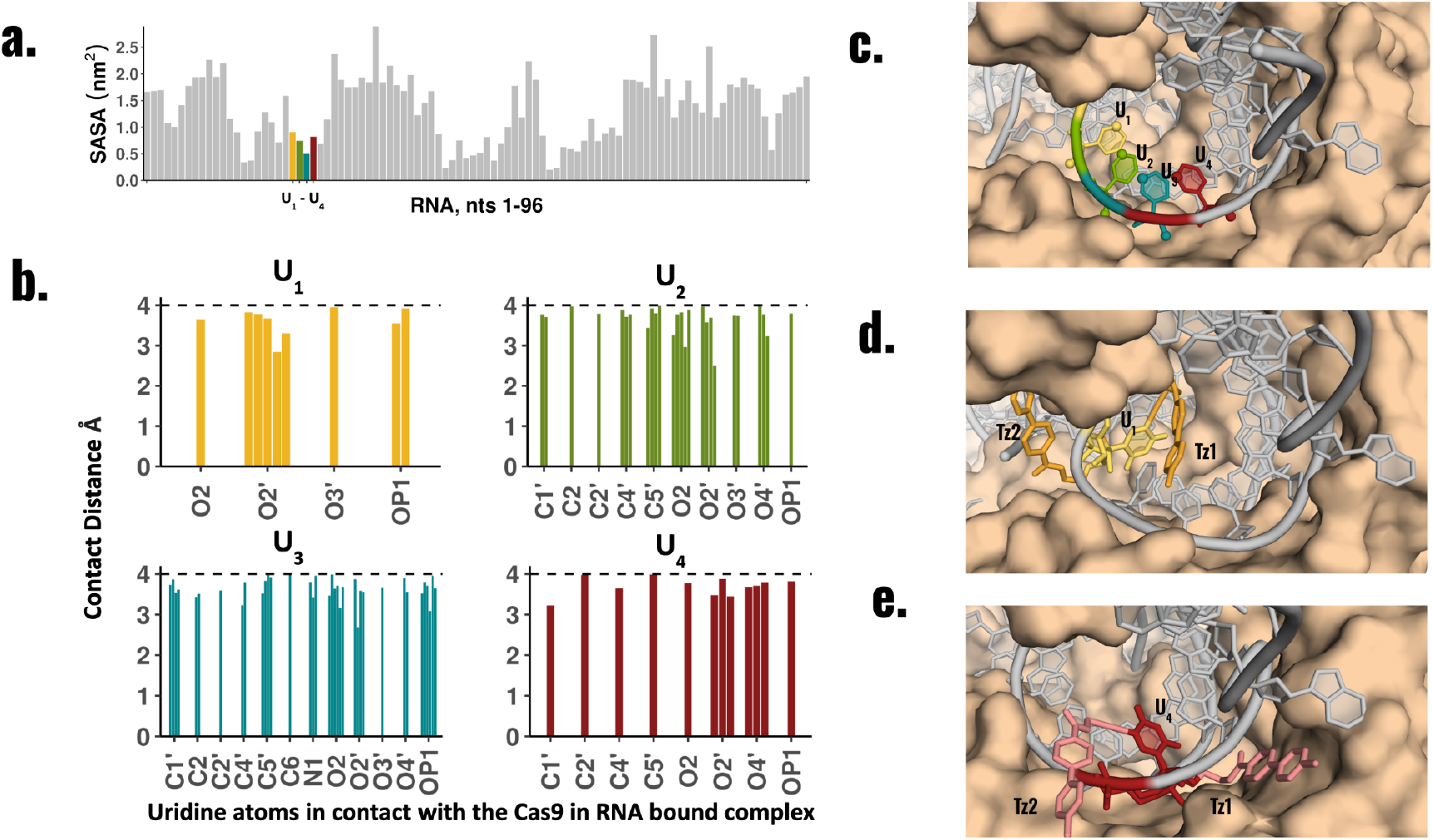
(**a**) Solvent accessible surface area of the RNA nucleotides in the protein:RNA:DNA complex (PDB ID: 4OO8) (**b**) Atomic contacts within 4Å of the four Us of the repeat sequence with Cas9, in the sgRNA-Cas9 complex. (**c**) Structural context of the four Us in the sgRNA-Cas9 complex. The potential modification sites (atoms C5 and O2’) are shown as spheres. sgRNA is shown in gray and Cas9 is shown in brown. (**d**) Tz_1_ and Tz_2_ modifications modeled on U_1_ (**e**) Tz_1_ and Tz_2_ modifications modeled on U_4_.

Following these predictions, we synthesized four sgRNAs targeting linearized pBR322 plasmid by solid phase RNA synthesis. The modified uridine phosphoramidites **1** and **2** (**Figure 2B**) were incorporated into positions U_1_ or U_4_. Chemical synthesis of these phosphoramidites has previously been reported.^24^ Tz was tagged post-synthetically using **Tz-NHS** (**Figure 2B**). We termed the constructs **sgRNA1-(U_1_Tz_1_), sgRNA1-(U_1_Tz_2_), sgRNA1-(U_4_Tz_1_)** and **sgRNA1-(U_4_Tz_2_)** to indicate the type of Tz modification and the position where it was installed. Following the synthesis, the four Tz-tagged sgRNAs were purified by preparative PAGE. Successful synthesis and purification were confirmed by denaturing gel electrophoresis and ESI-MS (**Figure S1**). We subsequently tested their ability to cut linearized pBR322 plasmid in the presence of Cas9. After 16 h of treatment the experiments were analyzed by agarose gel electrophoresis, shown in **Figure 4. sgRNA1-(U_4_Tz_1_)** was found to have the same level of activity as the unmodified **sgRNA1. sgRNA1-(U_4_Tz_2_)** was also found to have strong activity that was slightly lower than the unmodified **sgRNA1**. Meanwhile, modification at U_1_-position with either Tz_1_ or Tz_2_ significantly decreased nuclease activity.

**Figure 4.**
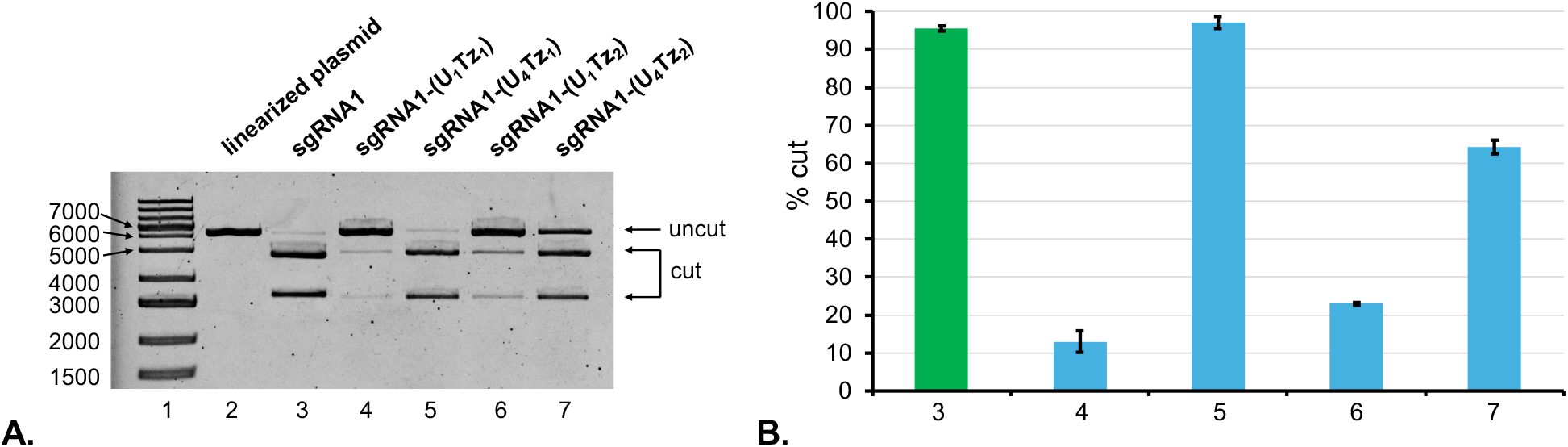
Analysis of CRISPR-Cas9 experiments using agarose gel electrophoresis. (**A**) *Lane 1*: DNA ladder; *Lane 2*: linearized pBR322 plasmid; *Lane 3*: linearized pBR322 plasmid and **sgRNA1**; *Lane 4*: linearized pBR322 plasmid and **sgRNA1-(U_1_Tz_1_)**; *Lane 5*: linearized pBR322 plasmid and **sgRNA1-(U_4_Tz_1_)**; *Lane 6*: linearized pBR322 plasmid and **sgRNA1-(U_1_Tz_2_)**; *Lane 7*: linearized pBR322 plasmid and **sgRNA1-(U_4_Tz_2_)**. (**B**) Bar chart representing the percentage of cut DNA upon treatment with the constructs in part **A**. Lane numbering is the same as in part **A**. All CRISPR experiments were performed in duplicate. Error bars represent ± s.d.

To test the generality of our tagging approach, we synthesized sgRNAs targeting linearized eGFP-N1 plasmid. We analogously termed them **sgRNA2-(U_1_Tz_1_), sgRNA2-(U_1_Tz_2_), sgRNA2-(U_4_Tz_1_)** and **sgRNA2-(U_4_Tz_2_)**. Following the synthesis, the four Tz-tagged sgRNAs were purified by preparative PAGE. and subsequently tested for their ability to cut linearized eGFP-N1 plasmid in the presence of Cas9. After 16 h of treatment the experiments were analyzed by agarose gel electrophoresis, shown in **Figure 5**. Once again, we observed that tagging of U_4_ with either Tz_1_ or Tz_2_ conserved the nuclease activity. Meanwhile, modification at U_1_-position with either Tz_1_ or Tz_2_ significantly lowered nuclease activity. The data indicate a general principle that sgRNAs tagged with either Tz_1_ or Tz_2_ at the U_4_ position of the repeat region will be constitutively active.

**Figure 5.**
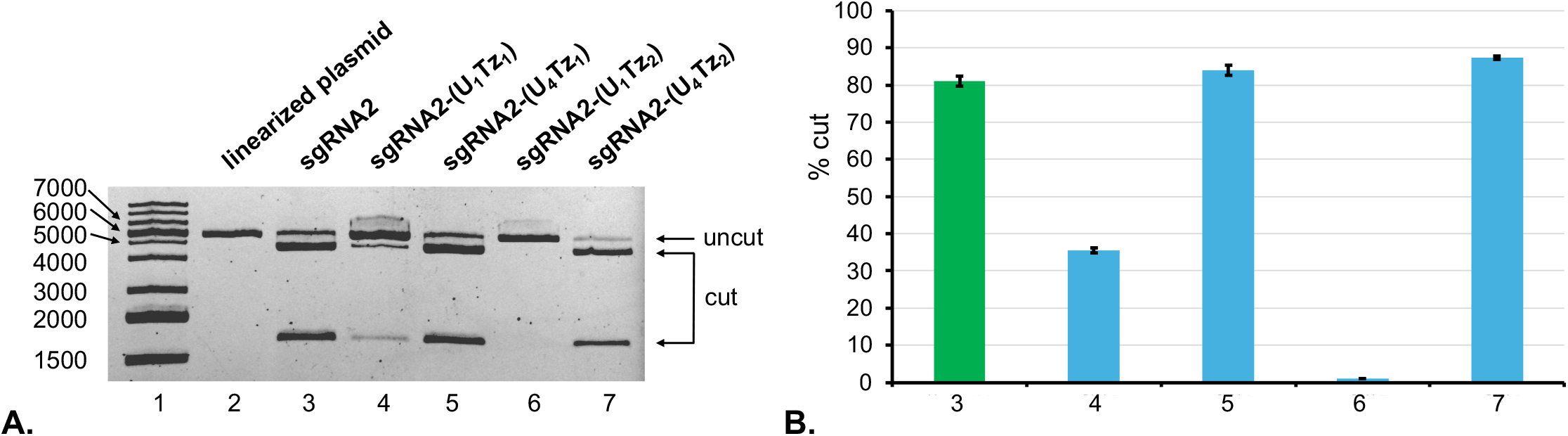
Analysis of CRISPR-Cas9 experiments using agarose gel electrophoresis. (**A**) *Lane 1*: DNA ladder; *Lane 2*: linearized eGFP-N1 plasmid; *Lane 3*: linearized eGFP-N1 plasmid and **sgRNA2**; *Lane 4*: linearized eGFP-N1 plasmid and **sgRNA2-(U_1_Tz_1_)**; *Lane 5*: linearized eGFP-N1 plasmid and **sgRNA2-(U_4_Tz_1_)**; *Lane 6*: linearized eGFP-N1 plasmid and **sgRNA2-(U_1_Tz_2_)**; *Lane 7*: linearized eGFP-N1 plasmid and **sgRNA2-(U_4_Tz_2_)**. (**B**) Bar chart representing the percentage of cut DNA upon treatment with the constructs in part **A**. Lane numbering is the same as in part **A**. All CRISPR experiments were performed in duplicate. Error bars represent ± s.d.

### (B) Design and synthesis of TCO-CRISPR suppressors

After achieving constitutively functional, Tz-tagged sgRNA, we synthesized four TCO-modified CRISPR suppressors, shown in **Figure 6**. The compounds have been designed based on 4 principles. They are modular, thus allowing for structural optimization and modification with TCO. They contain functional groups that will interfere with structural elements of sgRNA/Cas9 complex. They are based on the classes of compounds with known cell permeability. They are small molecules with molecular weights under 1200 Da, which is also important to facilitate cell permeability.

**Figure 6.**
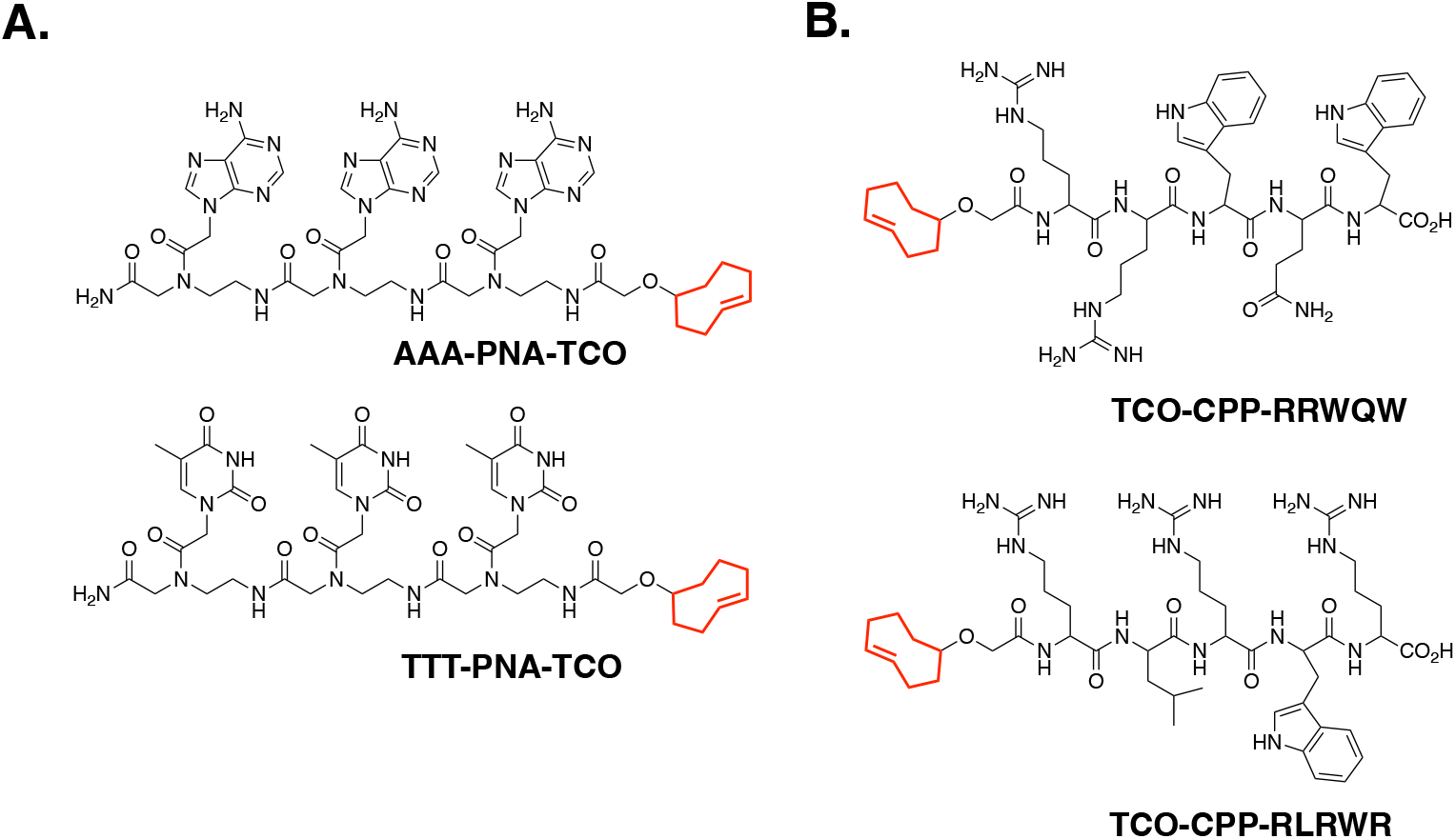
A library of small molecule TCO-modified CRISPR suppressors that are based on (**A**.) PNA; (**B**.) CPP.

The first set of suppressor compounds (**Figure 6A**) are peptide nucleic acids (PNAs) containing either thymine or adenine nucleobases. Although long PNA molecules are known to have poor cell permeability, short PNAs have been reported to be cell permeable.^25^ We hypothesize that either thymine or adenine nucleobases of the PNAs will interfere with the native repeat-anti-repeat structure that contains five U-A base pairs. Because PNAs can be synthesized in a programmable manner by solid phase synthesis they can be further optimized in terms of length and sequence to find the optimal disrupter of sgRNA/Cas9 complex.

The second set of compounds are TCO-CPP constructs (**Figure 6B**) that are based on the smallest-known cell-penetrating peptides (CPPs).^26–28^ They are also small molecules with MW under 1200 Da. As the name implies, they have been shown to be cell permeable and capable of shuttling various therapeutic cargoes across the cellular membrane. These short peptides can be easily synthesized by standard solid phase peptide synthesis. Post-synthetically, they were modified with TCO and purified by HPLC. In addition to inherent steric hinderance, guanidinium groups of the shown TCO-CPP constructs were expected to interfere with nucleobase base-pairing in the repeat:anti-repeat region of sgRNA.

#### TCO-CPP-RLRWR

We investigated the impact of TCO-modified CRISPR suppressors on the Cas9-assisted nuclease activity of **sgRNA1-(U_4_-Tz_1_)** and **sgRNA1-(U_4_Tz_2_)**. The pBR322 plasmid was treated with Cas9 and **sgRNA1-(U_4_-Tz_1_)** and **sgRNA1-(U_4_Tz_2_)** alone or in the presence of TCO-modified CRISPR suppressors. After 16 h of treatment the experiments were analyzed by agarose gel electrophoresis, shown in **Figure 7A**. The gel data was quantitated using ImageJ and plotted as a bar graph in **Figure 7B. TCO-CPP-RRWQW** and **TCO-CPP-RLRWR** completely inhibited activity of **sgRNA1-(U_4_Tz_2_)** (lanes 3 and 4). However, **sgRNA1-(U_4_Tz_2_)** treated with **TTT-PNA-TCO** or **AAA-PNA-TCO** still retained some residual activity (lanes 5 and 6). In the case of **sgRNA1-(U_4_Tz_1_)**, addition of **TTT-PNA-TCO** resulted in a residual nuclease activity (lane 10). **TCO-CPP-RRWQW, TCO-CPP-RLRWR** and **AAA-PNA-TCO** completely inhibited nuclease activity.

**Figure 7.**
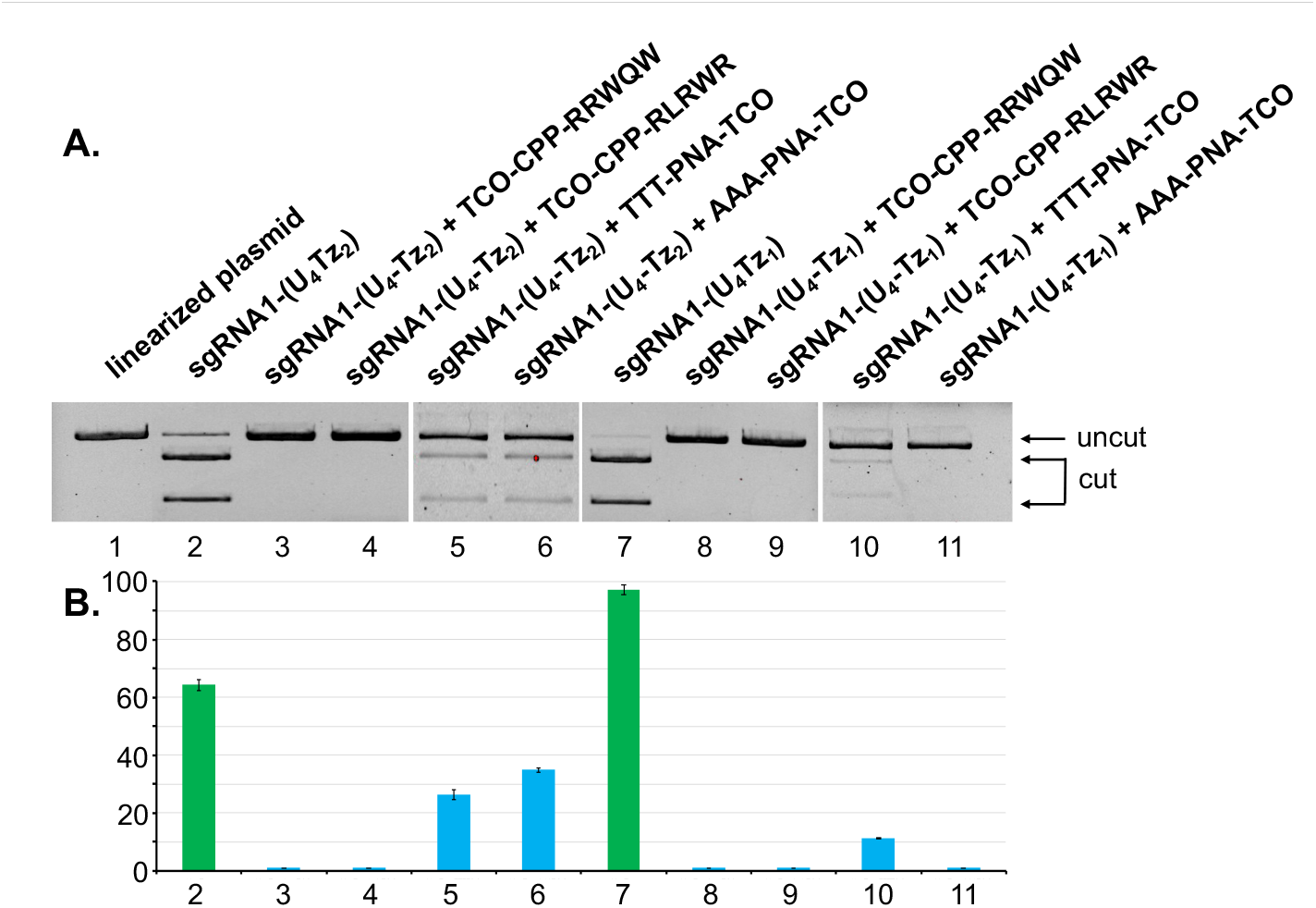
Analysis of CRISPR-Cas9 experiments in the presence of TCO-modified CRISPR suppressors. (**A**) *Lane 1*: linearized pBR322 plasmid; *Lane 2*: linearized pBR322 plasmid and **sgRNA1-(U_4_Tz_2_)**; *Lane 3*: linearized pBR322 plasmid**, sgRNA1-(U_4_Tz_2_)** and **TCO-CPP-RRWQW**; *Lane 4*: linearized pBR322 plasmid**, sgRNA1-(U_4_Tz_2_)** and **TCO-CPP-RLRWR**; *Lane 5*: linearized pBR322 plasmid**, sgRNA1-(U_4_Tz_2_)** and **TTT-PNA-TCO**; *Lane 6*: linearized pBR322 plasmid**, sgRNA1-(U_4_Tz_2_)** and **AAA-PNA-TCO**; *Lane 7*: linearized pBR322 plasmid and **sgRNA1-(U_4_Tz_1_)**; *Lane 8*: linearized pBR322 plasmid**, sgRNA1-(U_4_Tz_1_)** and **TCO-CPP-RRWQW**; *Lane 9*: linearized pBR322 plasmid**, sgRNA1-(U_4_Tz_1_)** and **TCO-CPP-RLRWR**; *Lane 10*: linearized pBR322 plasmid**, sgRNA1-(U_4_Tz_1_)** and **TTT-PNA-TCO**; *Lane 11*: linearized pBR322 plasmid**, sgRNA1-(U_4_Tz_1_)** and **AAA-PNA-TCO**. (**B**) Bar chart representing the percentage of cut DNA upon treatment with the constructs in part **A**. Lane numbering is the same as in part **A**. All CRISPR experiments were performed in duplicate. Error bars represent ± s.d.

To test the generality of our CRISPR-Cas9 regulation approach, we performed analogous experiments using **sgRNA2-(U_4_-Tz_1_)** and **sgRNA2-(U_4_Tz_2_)** targeting linearized eGFP-N1 plasmid. After 16 h of treatment the experiments were analyzed by agarose gel electrophoresis, shown in **Figure 8A**. The gel data was quantitated using ImageJ and plotted as a bar graph in **Figure 8B**. In the case of **sgRNA2-(U_4_Tz_2_), TCO-CPP-RRWQW** and **TTT-PNA-TCO** addition resulted in a residual nuclease activity (lanes 3 and 5). Addition of **TCO-CPP-RLRWR** and **AAA-PNA-TCO** completely inhibited nuclease activity (lanes 4 and 6). In the case of **sgRNA2-(U_4_Tz_1_), TTT-PNA-TCO** partially lowered nuclease activity (lane 10). Meanwhile, **TCO-CPP-RRWQW, TCO-CPP-RLRWR** and **AAA-PNA-TCO** completely inhibited nuclease activity (lanes 8, 9 and 11). In conclusion, the *in vitro* data suggests that temporal control of CRISPR-Cas9 activity, schematically shown in **Figure 1**, can be achieved using sgRNA modified with either Tz_1_ or Tz_2_ at the U_4_ position of the repeat region. Inactivation of Tz-tagged sgRNAs can be best achieved using either **TCO-CPP-RRWQW** or **TCO-CPP-RLRWR**.

**Figure 8.**
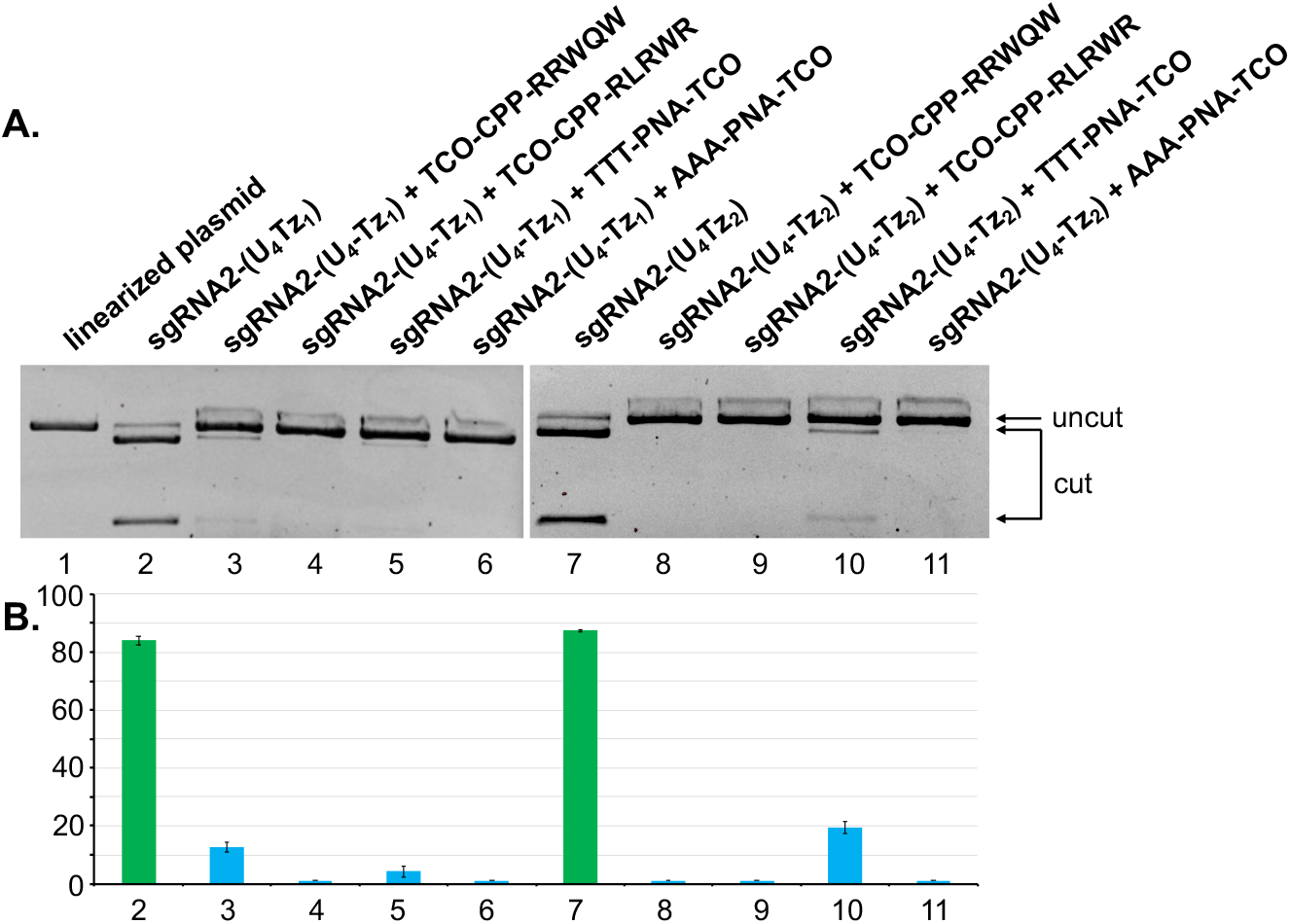
Analysis of CRISPR-Cas9 experiments in the presence of TCO-modified CRISPR suppressors. (**A**) *Lane 1*: linearized eGFP-N1 plasmid; *Lane 2*: linearized eGFP-N1 plasmid and **sgRNA2-(U_4_Tz_1_)**; *Lane 3*: linearized eGFP-N1 plasmid**, sgRNA2-(U_4_Tz_1_)** and **TCO-CPP-RRWQW**; *Lane 4*: linearized eGFP-N1 plasmid**, sgRNA2-(U_4_Tz_1_)** and **TCO-CPP-RLRWR**; *Lane 5*: linearized eGFP-N1 plasmid**, sgRNA2-(U_4_Tz_1_)** and **TTT-PNA-TCO**; *Lane 6*: linearized eGFP-N1 plasmid**, sgRNA2-(U_4_Tz_1_)** and **AAA-PNA-TCO**; *Lane 7*: linearized eGFP-N1 plasmid and **sgRNA2-(U_4_Tz_2_)**; *Lane 8*: linearized eGFP-N1 plasmid**, sgRNA2-(U_4_Tz_2_)** and **TCO-CPP-RRWQW**; *Lane 9*: linearized eGFP-N1 plasmid**, sgRNA2-(U_4_Tz_2_)** and **TCO-CPP-RLRWR**; *Lane 10*: linearized eGFP-N1 plasmid**, sgRNA2-(U_4_Tz_2_)** and **TTT-PNA-TCO**; *Lane 11*: linearized eGFP-N1 plasmid**, sgRNA2-(U_4_Tz_2_)** and **AAA-PNA-TCO**. (**B**) Bar chart representing the percentage of cut DNA upon treatment with the constructs in part **A**. Lane numbering is the same as in part **A**. All CRISPR experiments were performed in duplicate. Error bars represent ± s.d.

### C. Cell studies

We assessed cell permeability of the TCO-modified CRISPR suppressors using **OG-Tz** fluorescent probe.^29^ It’s a reported dye whose fluorescence is quenched by Tz. Fluorescence can be restored upon the click reaction with TCO. We tested fluorescence response of **OG-Tz** upon addition of the TCO-modified CRISPR suppressors in PBS (pH 7.4). As illustrated in **Figure S7**, there is a 10-fold enhancement of fluorescence. The cells were treated with **AAA-PNA-TCO, TTT-PNA-TCO, TCO-CPP-RRWQW** or **TCO-CPP-RLRWR** for 3 h. Afterwards, the cells were treated with **OG-Tz** (50 µM) for 2 h. Hoechst 33258 dye was used for nuclear staining. Cellular fluorescence was analyzed using microscopy and flow cytometry.

The microscopy data is shown in **Figure 9**. As expected, the cells treated with **OG-Tz** alone showed week fluorescence in the Oregon Green channel (**Figure 9C**). Punctate staining was observed in cells treated with **Tz-OG** and **AAA-PNA-TCO** and **TTT-PNA-TCO**, (**Figures 9G** and **9K**). The strongest fluorescence was observed in cells treated with **OG-Tz** and either **TCO-CPP-RRWQW** or **TCO-CPP-RLRWR** (**Figure 9O** and **9S**), suggesting that the CPP compounds have good cell permeability. The cells shown in **Figure 9** were trypsinized and analyzed by flow cytometry, which confirmed the findings (**Figure S8**).

**Figure 9.**
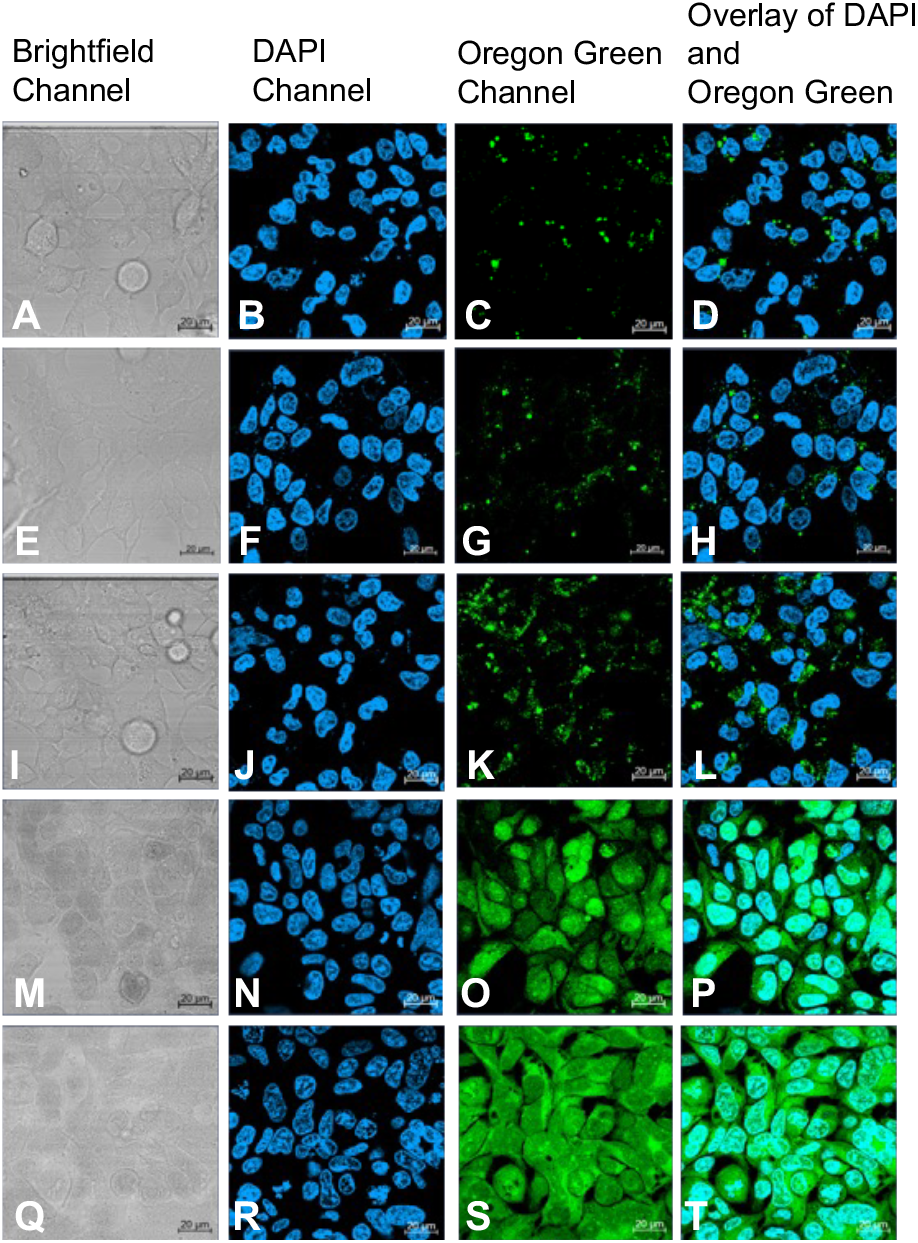
Assessment of cell permeability of TCO-modified CRISPR suppressors. HEK293 cells were treated with **OG-Tz** alone (A-D), or in combination with **AAA-PNA-TCO** (E-H), **TTT-PNA-TCO** (I-L), **TCO-CPP-RRWQW** (M-P) and **TCO-CPP-RLRWR** (Q-T).

Temporal control of CRISPR-Cas9 activity was assessed in GFP-expressing HEK239 cells. We were concerned about post-transfection stability of **sgRNA2** that might be exposed to nuclease degradation inside the cells. This concern was addressed by site-specific modification of **sgRNA2** with 2′-OMe groups. We followed the strategy described by Yin *et al.* who identified the exact positions which can be modified with 2′-OMe without significant perturbation to native binding between sgRNA and Cas9.^30^ We synthesized **sgRNA3**, having the sequence: 5′-GGGCGAGGAGCUGUUCACCGGU_1_U_2_U_3_U_4_AGagcuagaaauagcaaGUUaAaAuAaggcuaGUccG UUAucAAcuugaaaaagugGcaccgagucggugcuuuuu-3′ Capital letters indicate unmodified nucleotides, while small letters correspond to nucleotides containing 2′-OMe groups. For reference, uridines of the repeat region are numbered 1-4. U_4_ was tagged with either Tz_1_ or Tz_2_, thus making the constructs **sgRNA3-(U_4_Tz_1_)** and **sgRNA3-(U_4_Tz_2_)**. We assessed the CRISPR-Cas9 activity of these constructs *in vitro* (**Figures S9**). The *in vitro* experiments showed that 2′-OMe modifications did not perturb nuclease activity and the new constructs behave similarly to **sgRNA2-(U_4_Tz_1_)** and **sgRNA2-(U_4_Tz_2_)**.

GFP-expressing HEK239 cells were co-transfected with **sgRNA3-(U_4_Tz_1_)** or **sgRNA3-(U_4_Tz_2_)** and commercially available mRNA that encodes the Cas9 gene for 72 h. After the transfection, the media was replaced with fresh DMEM and the cells were allowed to grow for additional 48 h. GFP expression was analyzed by flow cytometry and compared to the untreated cells. As illustrated in **Figures 9E**, the mean fluorescence intensity (MFI) of GFP decreased by 44% after **sgRNA3-(U_4_Tz_1_)** treatment. Similarly, MFI of GFP decreased by 47% after **sgRNA3-(U_4_Tz_2_)** treatment (**Figure 10E**). Ability of TCO-modified CRISPR suppressors to control nuclease activity was examined in the next set of experiments. GFP-expressing HEK239 cells were co-transfected with **sgRNA3-(U_4_Tz_1_)** or **sgRNA3-(U_4_Tz_2_)** and commercially available mRNA that encodes the Cas9 gene for 48 h. Then, **AAA-PNA-TCO, TTT-PNA-TCO, TCO-CPP-RRWQW** and **TCO-CPP-RLRWR** (10 µM) were added to the media. After 24 h of treatment with the TCO-modified CRISPR suppressors, the media was replaced with fresh DMEM and the cells were allowed to grow for additional 48 h. As illustrated in **Figures 10A, 10E, 11A** and **11E, TCO-CPP-RRWQW** was able to inactivate **sgRNA3-(U_4_Tz_1_)** and **sgRNA3-(U_4_Tz_2_)** inside the cells. In these experiments, the MFI of GFP decreased by lower amounts as the result of turned off CRISPR-Cas9 complex. Analogous results were observed after treatment with **TCO-CPP-RLRWR** (**Figures 10B, 10E, 11B** and **11E**). However, the PNA-based CRISPR suppressors were not as effective, as shown in **Figures 10C, 10D, 10E, 11C, 11D** and **11E**. The observed MFI of GFP was very close to the cells treated with **sgRNA3-(U_4_Tz_1_)** or **sgRNA3-(U_4_Tz_2_)** alone. This is probably due to inferior cell permeability of **AAA-PNA-TCO** and **TTT-PNA-TCO**, as was observed in **Figure 9**.

**Figure 10.**
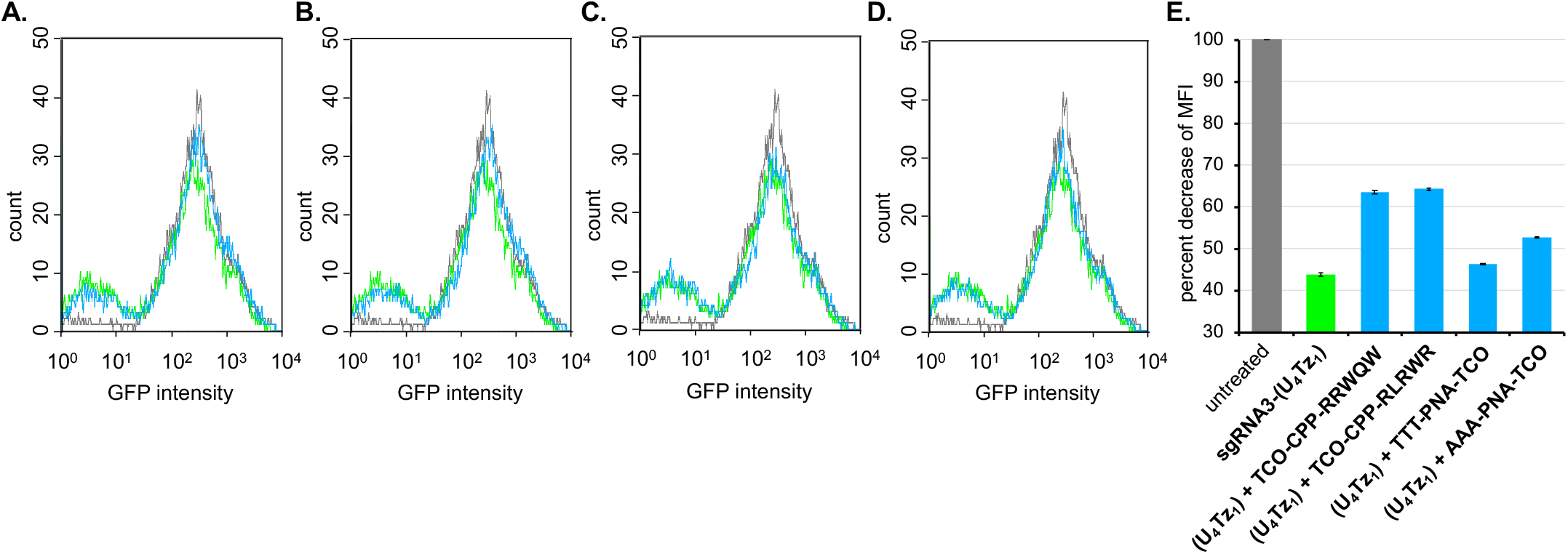
Flow cytometry of GFP-expressing HEK293 cells treated with **sgRNA 3-(U_4_Tz_1_)** and TCO-modified CRISPR suppressors. Histograms of untreated cells are shown in black. The cells treated with **sgRNA 3-(U_4_Tz_1_)** are shown in green. The cells treated with **sgRNA 3-(U_4_Tz_1_)** and TCO-modified CRISPR suppressors are shown in blue. (**A.**) overlay of histograms of the untreated cells, cells treated with **sgRNA 3-(U_4_Tz_1_)** and cells treated with **sgRNA 3-(U_4_Tz_1_)** and **TCO-CPP-RRWQW**; (**B.**) overlay of histograms of the untreated cells, cells treated with **sgRNA 3-(U_4_Tz_1_)** and cells treated with **sgRNA 3-(U_4_Tz_1_)** and **TCO-CPP-RLRWR**; (**C.**) overlay of histograms of the untreated cells, cells treated with **sgRNA 3-(U_4_Tz_1_)** and cells treated with **sgRNA 3-(U_4_Tz_1_)** and **TTT-PNA-TCO**; (**D.**) overlay of histograms of the untreated cells, cells treated with **sgRNA 3-(U_4_Tz_1_)** and cells treated with **sgRNA 3-(U_4_Tz_1_)** and **AAA-PNA-TCO**; (**E.**) decrease of MFI of GFP relative to the untreated GFP-expressing HEK293 cells. All experiments were performed in duplicate. Error bars represent ± s.d.

**Figure 11.**
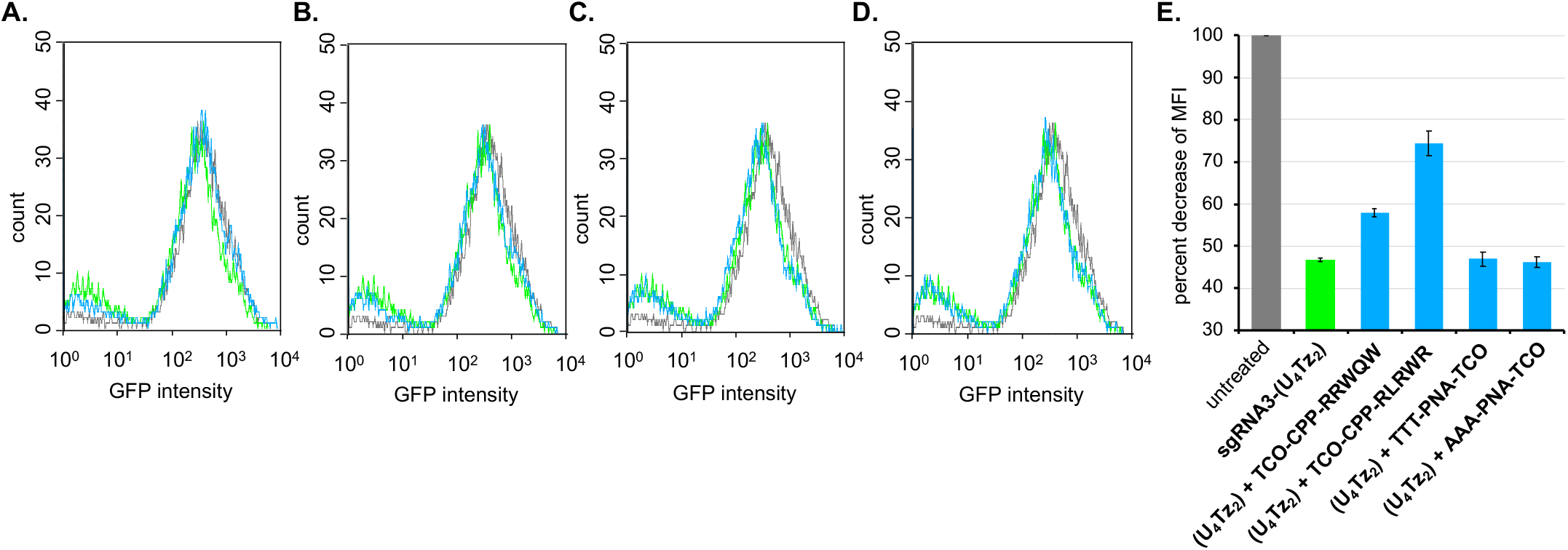
Flow cytometry of GFP-expressing HEK293 cells treated with **sgRNA 3-(U_4_Tz_2_)** and TCO-modified CRISPR suppressors. Histograms of untreated cells are shown in black. The cells treated with **sgRNA 3-(U_4_Tz_2_)** are shown in green. The cells treated with **sgRNA 3-(U_4_Tz_2_)** and TCO-modified CRISPR suppressors are shown in blue. (**A.**) overlay of histograms of the untreated cells, cells treated with **sgRNA 3-(U_4_Tz_2_)** and cells treated with **sgRNA 3-(U_4_Tz_2_)** and **TCO-CPP-RRWQW**; (**B.**) overlay of histograms of the untreated cells, cells treated with **sgRNA 3-(U_4_Tz_2_)** and cells treated with **sgRNA 3-(U_4_Tz_2_)** and **TCO-CPP-RLRWR**; (**C.**) overlay of histograms of the untreated cells, cells treated with **sgRNA 3-(U_4_Tz_2_)** and cells treated with **sgRNA 3-(U_4_Tz_2_)** and **TTT-PNA-TCO**; (**D.**) overlay of histograms of the untreated cells, cells treated with **sgRNA 3-(U_4_Tz_2_)** and cells treated with **sgRNA 3-(U_4_Tz_2_)** and **AAA-PNA-TCO**; (**E.**) decrease of MFI of GFP relative to the untreated GFP-expressing HEK293 cells. All experiments were performed in duplicate. Error bars represent ± s.d.

To illustrate the therapeutic potential of our technology, we implemented it towards a well-established medicinal target, vascular endothelial growth factor A (VEGFA). VEGFA is an angiogenic factor, whose expression is upregulated in neovascular age-related macular degeneration, the leading cause of vision loss.^31,32^ CRISPR-Cas9-based disruption of VEGFA has already been explored as a therapeutic strategy.^33,34^ There are also reported off-target effects associated with a CRISPR-Cas9 system targeting VEGFA. Towards achieving temporal control over CRISPR-Cas9 treatment, we designed **sgRNA4-(U_4_Tz_1_)** and **sgRNA4-(U_4_Tz_2_)** where the guide region was programmed to target the VEGFA gene.

To assess the new constructs, HEK239T cells were co-transfected with the unmodified **sgRNA4, sgRNA4-(U_4_Tz_1_)**, or **sgRNA4-(U_4_Tz_2_)** and commercially available mRNA that encodes the Cas9 gene for 72 h. After the transfection, the media was replaced with fresh DMEM and the cells were allowed to grow for additional 48 h. VEGFA expression was analyzed by Western blot shown in **Figure 12**. We observed marked decrease in VEGFA expression in sgRNA-Cas9 treated cells. The Tz-tagged sgRNAs had similar levels of VEGFA suppression as the unmodified **sgRNA4**. We repeated these experiments, while adding **TCO-CPP-RRWQW** and **TCO-CPP-RLRWR** (10 µM) after 48 hours of transfection. VEGFA expression was once again analyzed by Western blot. Addition of TCO-modified CRISPR suppressors resulted in higher VEGFA expression, suggesting that Tz-tagged sgRNAs were inactivated inside the cells.

**Figure 12.**
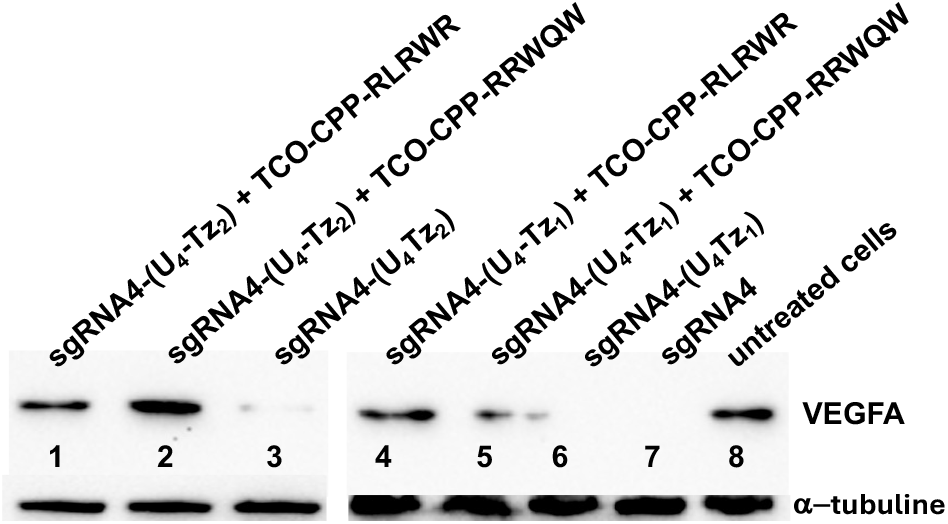
Western blot analysis of CRISPR-Cas9 experiments targeting VEGFA. Cells were treated with: *Lane 1*: **sgRNA4-(U_4_Tz_2_)** and **TCO-CPP-RLRWR**; *Lane 2*: **sgRNA4-(U_4_Tz_2_)** and **TCO-CPP-RRWQW**; *Lane 3*: **sgRNA4-(U_4_Tz_2_)**; *Lane 4*: linearized eGFP-N1 plasmid**, sgRNA2-(U_4_Tz_1_)** and **TCO-CPP-RLRWR**; *Lane 5*: **sgRNA4-(U_4_Tz_1_)** and **TCO-CPP-RLRWR**; *Lane 6*: **sgRNA4-(U_4_Tz_1_)** and **TCO-CPP-RRWQW**; *Lane 7*: **sgRNA4-(U_4_Tz_1_)**; *Lane 8*: untreated cells. α-tubulin was used as a loading control.

## Conclusion

Herein we described a system that facilitates temporal control of CRISPR-Cas9 nuclease activity using bio-orthogonal chemistry. We identified a precise position within the repeat:anti-repeat region of sgRNA that can be tagged with Tz without disrupting Cas9-enabled nuclease activity. We carried out Tz-tagging of three different sgRNAs to illustrate that this strategy is general. Afterwards, we identified TCO-modified small molecule CRISPR suppressors that click to Tz and significantly reduce the nuclease activity. Of the tested molecules, CPP-based suppressors showed the most effective reduction of CRISPR activity both in solution and live cells. Overall, the described strategy to control CRISPR-Cas9 nuclease activity consists of a single small-molecule tag and small molecule cell-permeable suppressor. Lastly, we illustrated the therapeutic potential of our technology by targeting the VEGFA gene in live HEK293T cells. In the future, we plan to examine if our technology for temporal control of CRISPR-Cas9 nuclease activity can reduce off-target effects using GUIDE-Seq protocol.^8^ We also plan to expand the library of TCO-modified small molecule CRISPR suppressors to investigate the precise mechanism of inactivation of nuclease activity.

