## Supporting Information for "Temporal Control of CRISPR-Cas9 Activity Using Bio-orthogonal Chemistry"


| **Table of Contents** | **Figure and Scheme titles** | **Page** |
|---|---|---|
| Materials and Methods | | S2-S6 |
| Figure S1 | PAGE analysis of purified sgRNAs | S7 |
| Figure S2 | PAGE analysis of purified sgRNAs | S7 |
| Figure S3 | Analytical HPLC and ESI-MS spectra of **TCO-CPP-RRWQW** | S8 |
| Figure S4 | Analytical HPLC and ESI-MS spectra of **TCO-CPP-RLRWR** | S9 |
| Figure S5 | Analytical HPLC and ESI-MS spectra of **AAA-PNA-TCO** | S10 |
| Figure S6 | Analytical HPLC and ESI-MS spectra of **TTT-PNA-TCO** | S11 |
| Figure S7 | Fluorescence spectra of **OG-Tz** conjugated to different TCO-modified CRISPR suppressors | S12 |
| Figure S8 | Flow cytometry of HEK293 cells treated with **OG-Tz** alone and with **OG-Tz** and TCO-modified CRISPR suppressors | S12 |
| Figure S9 | Analysis of CRISPR-Cas9 experiments using agarose gel electrophoresis | S13 |

### MATERIALS AND METHODS

All oligonucleotide solid phase syntheses were done on a 1.0 μmol scale using the Oligo-800 synthesizer (Azco Biotech, Oceanside, CA, USA). Solid phase syntheses were performed on control-pore glass (CPG-1000) purchased from Glen Research (Sterling, VA, USA). Other oligonucleotide solid phase synthesis reagents were obtained from ChemGenes Corporation (Wilmington, MA, USA). Phosphoramidites (TBDMS as the 2′-OH protecting group): rA was N-Bz protected, rC was N-Ac protected and rG was N-iBu protected. A, C, G, U phosphoramidites were dissolved in anhydrous acetonitrile (0.07 M) directly before use. $m^1A$, $m^6A$, $s^2U$ and $s^4U$ phosphoramidites were dissolved in anhydrous acetonitrile (0.15 M) directly before use. Coupling step was done using 5-ethylthio-1H-tetrazole solution (0.25 M) in acetonitrile for 12 min. 5′-detritylation step was done using 3% trichloroacetic acid in $CH_2Cl_2$. Oxidation step was done using $I_2$ (0.02 M) in THF/pyridine/$H_2O$ solution.

For gel electrophoresis, 10X Tris/Borate/EDTA (TBE) buffer was purchased from Fisher Scientific Company L.L.C. (Waltham, MA, USA) and used with proper dilution. 30% Arcylamide/Bis-arcylamide solution (29:1) was purchased from Bio-Rad Laboratories, Inc. (Hercules, CA, USA). GeneRuler 1 kb Plus DNA Ladder (cat.# FERSM1331) was purchased from Fisher Scientific. Chromatographic purifications of synthetic materials were conducted using SiliaSphere™ spherical silica gel with an average particle and pore size of 5 μm and 60 Å, respectively (Silicycle Inc, QC, Canada). Thin layer chromatography (TLC) was performed on SiliaPlate™ silica gel TLC plates with 250 μm thickness (Silicycle Inc, QC, Canada). Flash chromatography was performed using Biotage Isolara One instrument (Biotage Sweden AB, Uppsala, Sweden). Preparative TLC was performed using SiliaPlate™ silica gel TLC plates with 1000 μm thickness. $^1H$, $^{13}C$ and $^{31}P$ NMR spectroscopy was performed on a Bruker NMR at 500 MHz ($^1H$) and 126 MHz ($^{13}C$). All $^{13}C$ NMR spectra were proton decoupled. High resolution ESI-MS spectra of small molecules was acquired using Agilent Technologies 6530 Q-TOF instrument.

The peptides, **RRWQW** and **RLRWR**, were purchased from GenScript. PNAs were synthesized using Fmoc-Rink Amide AM Resin (Aapptec, cat. # RRZ001). PNA monomers, Fmoc-PNA-T-OH and Fmoc-PNA-A(Bhoc)-OH), were purchased from Biosearch Technologies (cat. # LK5004-B500 and LK5001-B500). PNAs were synthesized using the procedure described by Braasch, D. A. *et al. Current Protocols in Nucleic Acid Chemistry*, **2002**, 4.11.1-4.11.18.

**RNA Sequences:**

**pBR322-targeting sgRNA:**

**sgRNA1-($U_1Tz_1$)**
5′ - GGGCGCUUGUUUCGGCGUGGGUAG**$U_1$-Tz1$U_2U_3U_4$**AGAGCUAGACAUAGC
AAGUUAAAAUAAGGCUAGUCCGUUAUCAACUUGAAAAAGUGGCACCGAGUCGGU
GCUUUU - 3′

**sgRNA1-($U_4Tz_1$)**
5′ -GGGCGCUUGUUUCGGCGUGGGUAG**$U_1U_2$U3$U_4$-Tz1**AGAGCUAGACAUAGC
AAGUUAAAAUAAGGCUAGUCCGUUAUCAACUUGAAAAAGUGGCACCGAGUCGGU
GCUUUU - 3′

**sgRNA1-($U_1Tz_2$)**
5′ - GGGCGCUUGUUUCGGCGUGGGUAG**$U_1$-Tz2$U_2U_3U_4$**AGAGCUAGACAUAGC
AAGUUAAAAUAAGGCUAGUCCGUUAUCAACUUGAAAAAGUGGCACCGAGUCGGU
GCUUUU - 3′

**sgRNA1-($U_4Tz_2$)**:
5′ -GGGCGCUUGUUUCGGCGUGGGUAG**$U_1U_2$U3$U_4$-Tz2**AGAGCUAGACAUAGC
AAGUUAAAAUAAGGCUAGUCCGUUAUCAACUUGAAAAAGUGGCACCGAGUCGGU
GCUUUU - 3′

**eGFP-targeting sgRNA:**

**sgRNA2-($U_1Tz_1$)**
5'-GGGCGAGGAGCUGUUCACCGG**$U_1$-Tz1$U_2U_3U_4$**AGAGCUAGAAAUAGCAAGUU
AAAAUAAGGCUAGUCCGUUAUCAACUUGAAAAAGUGGCACCGAGUCGGUGCUUU
UU-3'

**sgRNA2-($U_4Tz_1$)**
5'-GGGCGAGGAGCUGUUCACCGG**$U_1U_2U_3U_4$-Tz1**AGAGCUAGAAAUAGCAAGUU
AAAAUAAGGCUAGUCCGUUAUCAACUUGAAAAAGUGGCACCGAGUCGGUGCUUU
UU-3'

**sgRNA2-($U_1Tz_2$)**
5'-GGGCGAGGAGCUGUUCACCGG**$U_1$-Tz2$U_2U_3U_4$**AGAGCUAGAAAUAGCAAGUU
AAAAUAAGGCUAGUCCGUUAUCAACUUGAAAAAGUGGCACCGAGUCGGUGCUUU
UU-3'

**sgRNA2-($U_4Tz_2$)**
5'-GGGCGAGGAGCUGUUCACCGG**$U_1U_2U_3U_4$-Tz2**AGAGCUAGAAAUAGCAAGUU
AAAAUAAGGCUAGUCCGUUAUCAACUUGAAAAAGUGGCACCGAGUCGGUGCUUU
UU-3'

**eGFP-targeting sgRNA, containing 2′-OMe groups (small letters):**

**sgRNA3**
5′-GGGCGAGGAGCUGUUCACCGG**UUUU**AGagcuagaaauagcaaGUU
aAaAuAaggcuaGUccGUUAucAAcuugaaaaagugGcaccgagucggugcuuuuu-3′

**sgRNA3-($U_4Tz_1$)**
5′-GGGCGAGGAGCUGUUCACCGG**UUU$U_4$-Tz1**AGagcuagaaauagcaaGUU
aAaAuAaggcuaGUccGUUAucAAcuugaaaaagugGcaccgagucggugcuuuuu-3′

**sgRNA3-($U_4Tz_2$)**
5′-GGGCGAGGAGCUGUUCACCGG**UUU$U_4$-Tz2**AGagcuagaaauagcaaGUU
aAaAuAaggcuaGUccGUUAucAAcuugaaaaagugGcaccgagucggugcuuuuu-3′

**VEGFA-targeting sgRNA, containing 2′-OMe groups (small letters):**

**sgRNA4**
**5'-**GGUGAGUGAGUGUGUGCGUGG**UUU**AGagcuagaaauagcaaGUU
aAaAuAaggcuaGUccGUUAucAAcuugaaaaagugGcaccgagucggugcuuuuu-3′

**sgRNA4-($U_4$Tz$_1$)**
**5'-**GGUGAGUGAGUGUGUGCGUGG**UUU**$U_4$**-Tz1**AGagcuagaaauagcaaGUU
aAaAuAaggcuaGUccGUUAucAAcuugaaaaagugGcaccgagucggugcuuuuu-3′

**sgRNA4-($U_4$Tz$_2$)**
**5'-**GGUGAGUGAGUGUGUGCGUGG**UUU**$U_4$**-Tz2**AGagcuagaaauagcaaGUU
aAaAuAaggcuaGUccGUUAucAAcuugaaaaagugGcaccgagucggugcuuuuu-3′

Capital letters indicate unmodified nucleotides, while small letters correspond to nucleotides containing 2′-OMe groups.

**CRISPR-Cas9 in vitro DNA cleavage assay:**

eGFP-N1 plasmid DNA (10 U/μL, 1 μL, NEB, R3510L) was diluted with water (16.87 μL) and NEB buffer 3.1 (10x, 2 μL). The plasmid was linearized directly prior to CRISPR with DraIII-HF (10 U/μL, 1 μL, NEB, R3510L). For the Cas9-mediated DNA cleavage assay, sgRNA (300 nM, 5 μL), Cas9 (1 μM, 0.3 μL, NEB, M0386S), Cas9 buffer (10x, 1 μL, NEB), linearized plasmid (20 nM, 1.5 μL) and MQ $H_2O$ (2.2 μL) were mixed (final volume = 10 μL) and incubated for 16 h at 37 °C. CRISPR experiments were terminated by the addition of proteinase K (20 mg/mL, 0.5 μL) for 1 h at 37 °C. The reaction (10μL) was mixed with blue loading buffer (6x, 2 μL, NEB, B7703S) and loaded on a 1% agarose stained with ethidium bromide (1x TBE running buffer).

**CRISPR-Cas9 experiments in HEK293 cells:**

CRISPR-Cas9 experiments, were carried out following the procedure reported by Yin, H. et al. [Nat. Chem. Biol. 2018, 14, 311-316]. The GFP-expressing HEK293 cells were purchased from GenTarget (cat# SC001) and cultured in DMEM, containing 10% FBS and 1X Penicillin/Streptomycin, at 37 °C, 5% $CO_2$, and 95% humidity. The cells were seeded at a concentration of $1 \times 10^5$ cells per well in 6-well plate 24 h prior to the experiment. The cells were transfected with Cas9 mRNA (500 ng, Thermo Fisher Scientific), GFP-targeting **sgRNAs** (30 nM) using lipofectamine (1.5 μL) (Invitrogen™ LMRNA003) for 72 h in Opti-MEM reduced serum media. After 72 h, Opti-MEM was replaced with fresh DMEM and the cells were grown for additional 48 h. The cells were treated with trypsin for 5 min, collected by centrifugation at 1000 RPM and suspended in PBS (1 mL). GFP expression was analyzed by flow cytometry. Data from $10^6$ cells were acquired using a FACS Aria III cell sorter equipped with a 488 nm/blue coherent sapphire solid-state laser, 20 mW (BD Biosciences, San Jose, CA, USA). Data analyses were carried out using FlowJo software (Ashland, OR, USA), according to manufacturer's instructions. Parameters, such as MFI and the percentages of specific populations were quantified by histogram analysis.

Ability of TCO-modified CRISPR suppressors to control nuclease activity was examined as follows: GFP-expressing HEK239 cells were co-transfected with Cas9 mRNA (500 ng, Thermo

Fisher Scientific), GFP-targeting **sgRNAs** (30 nM) using lipofectamine (1.5 μL) (Invitrogen™ LMRNA003) for 48 h. Then, **AAA-PNA-TCO**, **TTT-PNA-TCO**, **TCO-CPP-RRWQW** and **TCO-CPP-RLRWR** (10 μM) were added to the media. After 24 h of treatment with the TCO-modified CRISPR suppressors, the media was replaced with fresh DMEM and the cells were allowed to grow for additional 48 h. The cells were treated with trypsin for 5 min, collected by centrifugation at 1000 RPM and suspended in PBS (1 mL). GFP expression was analyzed by flow cytometry. Data from $10^6$ cells were acquired using a FACS Aria III cell sorter equipped with a 488 nm/blue coherent sapphire solid-state laser, 20 mW (BD Biosciences, San Jose, CA, USA). Data analyses were carried out using FlowJo software (Ashland, OR, USA), according to manufacturer's instructions. Parameters, such as MFI and the percentages of specific populations were quantified by histogram analysis.

**Assessment of cell permeability of TCO-modified CRISPR suppressors.**

The HEK293 cells were cultured in DMEM, containing 10% FBS and 1X Penicillin/Streptomycin, at 37 °C, 5% $CO_2$, and 95% humidity. The cells were seeded at a concentration of $1 \times 10^5$ cells per well in 6-well plate 24 h prior to the experiment. The cells were treated with the TCO-modified CRISPR suppressors (30 μM) for 3 h. Then, the media was replaced and the cells were treated with **OG-Tz** (50 μM) for 2h. Afterwards, the cells were treated with trypsin for 5 min, collected by centrifugation at 1000 RPM and suspended in PBS (1 mL). GFP expression was analyzed by flow cytometry. Data from $10^6$ cells were acquired using a FACS Aria III cell sorter equipped with a 488 nm/blue coherent sapphire solid-state laser, 20 mW (BD Biosciences, San Jose, CA, USA). Data analyses were carried out using FlowJo software (Ashland, OR, USA), according to manufacturer's instructions. Parameters, such as MFI and the percentages of specific populations were quantified by histogram analysis.

HEK293 cells were grown to ~70% confluence in 35 mm MatTek glass bottom dishes in DMEM supplemented with 10% bovine serum albumin, penicillin and streptomycin. The cells were treated with **AAA-PNA-TCO**, **TTT-PNA-TCO**, **TCO-CPP-RRWQW** or **TCO-CPP-RLRWR** (30 μM) for 3 h. Afterwards, the cells were treated with **OG-Tz** (50 μM) for 2 h. Hoechst 33258 dye was used for nuclear staining. Cellular fluorescence was analyzed using confocal microscopy. Microscopy experiments were carried out using Zeiss LSM980 confocal microscope.

**Modeling Studies**

The structure of the Cas9 in complex with RNA and DNA was obtained from the Protein Data Bank (PDB ID : 4OO8 ; Nishimasu, H. *et al.* **2014**, *Cell*, *156*(5), 935 - 949). Solvent accessible surface area (SASA) and number and distance of atomic contacts were calculated for the RNA bound to Cas9 using Pymol (The PyMOL Molecular Graphics System, Version 3.0 Schrödinger, LLC). The Tz1 and Tz2 modifications were introduced on the RNA repeat sequence in MOE (Molecular Operating Environment (MOE), 2024.06 Chemical Computing Group ULC, 910-1010 Sherbrooke St. W., Montreal, QC H3A 2R7, 2024). Local minimization of the structure was then carried out to accommodate the modifications in the complex.

**Modification of sgRNAs with Tz**

sgRNA (263 μM) was dissolved in borate buffer (250 μL, pH = 9.4). Compound **Tz-NHS** was dissolved in DMF (400μL, 50 mM) and added to the RNA. The reaction mixture was vortexed and agitated at 1000 rpm at rt for 8 h. The conjugated sgRNA was purified from small molecules using Amicon Ultra 3K column (MilliporeSigma, cat# UFC500396) and washed with MQ $H_2O$ (3x

300 μL). sgRNAs were purified by preparative PAGE. Purified sgRNAs were analyzed using analytical PAGE, as shown in **Figure S1** and **Figure S2**.

**Western blot analysis**

Total protein lysate was harvested in RIPA buffer and proteins separated by 10% SDS-PAGE were transferred to a PVDF membrane by using High MW protocol on Biorad Turbo wet transfer. Following transfer, membranes were blocked in 5% milk in PBST for 1 h at room temperature, washed three times in PBS with 0.1% (w/v) Tween 20 (PBS/T) for 10 min each and placed in primary antibody overnight at 4 °C. Primary antibody was rabbit VEGFA polyclonal antibody (1:2000 dilution, Cat no:19003-1-AP, Proteintech) and α-tubulin monoclonal antibody (1:20,000 dilution, Cat no.666031-1-Ig, Proteintech) in 5% BSA in PBS/T. Following overnight incubation, membranes were washed three times in PBS/T for 10 min each. Then membranes were incubated in a 1:10,000 dilution of goat anti-rabbit secondary (Jackson), goat-anti mouse secondary in 5% milk and PBS/T. Membranes were washed a final 3 times in PBS/T for 10 min each prior to being imaged on the BioRad ChemiDoc by using Thermo Scientific SuperSignal West Pico PLUS Chemiluminescent Substrate (Catalog: 34577).

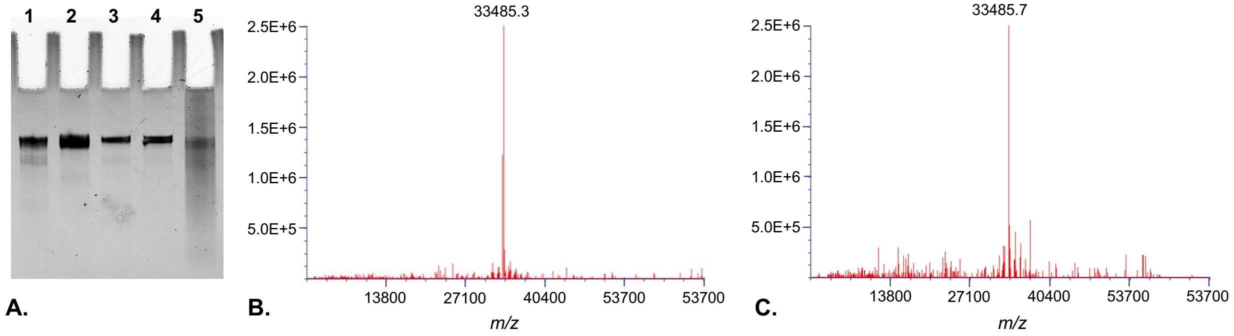


**Figure S1**. (**A.**) PAGE analysis of purified sgRNAs: *Lane 1*: **sgRNA1-(U$_4$Tz$_2$)**; *Lane 2*: **sgRNA1-(U$_1$Tz$_2$)**; *Lane 3*: **sgRNA1-(U$_4$Tz$_1$)**; *Lane 4*: **sgRNA1-(U$_1$Tz$_1$)**; *Lane 5*: reference. (**B.**) Deconvoluted ESI-MS analysis of **sgRNA1-(U$_1$Tz$_2$)**; (**C.**) Deconvoluted ESI-MS analysis of **sgRNA1-(U$_4$Tz$_2$)**.

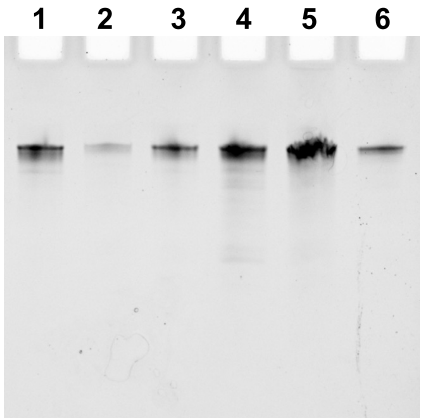


**Figure S2**. PAGE analysis of purified sgRNAs: *Lane 1*: **sgRNA2-(U$_4$Tz$_2$)**; *Lane 2*: **sgRNA2-(U$_1$Tz$_2$)**; *Lane 3*: **sgRNA2-(U$_4$Tz$_1$)**; *Lane 4*: **sgRNA2-(U$_1$Tz$_1$)**; *Lane 5*: unmodified **sgRNA2**; *Lane 6*: reference.

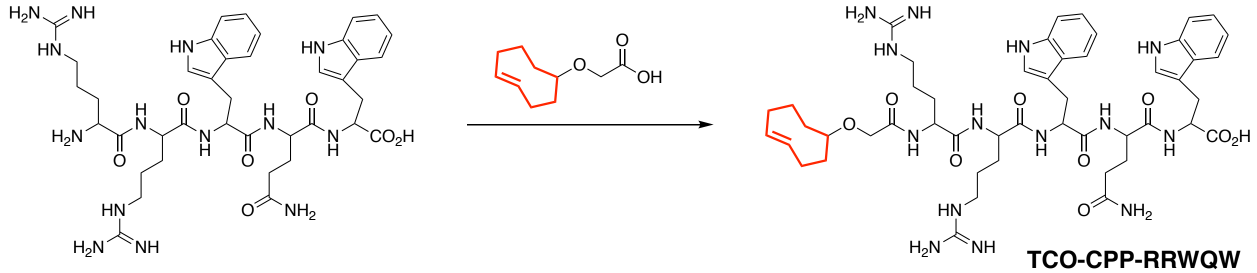


Prepared a suspension of HATU (14 mg) in DMF (200 μL). In parallel, prepared a solution of (*E*)-2-(Cyclooct-4-en-1-yloxy)acetic acid (3.3 mg) in NMP (100 μL). Combined the two solutions, added DIPEA (7μL) and 2,6-lutidine (7μL) and stirred for 10 min at rt. Added a solution of **RRWQW** (20 mg, 24 μmol) in DMF (100 μL) and stirred at rt for 2 h. **TCO-CPP-RRWQW** was purified by preparative HPLC and analyzed by analytical HPLC and ESI-MS, shown below. HRMS (ESI) cal'd for $C_{49}H_{69}N_{14}O_9$ $[M+1]^+$ 997.5366; observed 997.5381

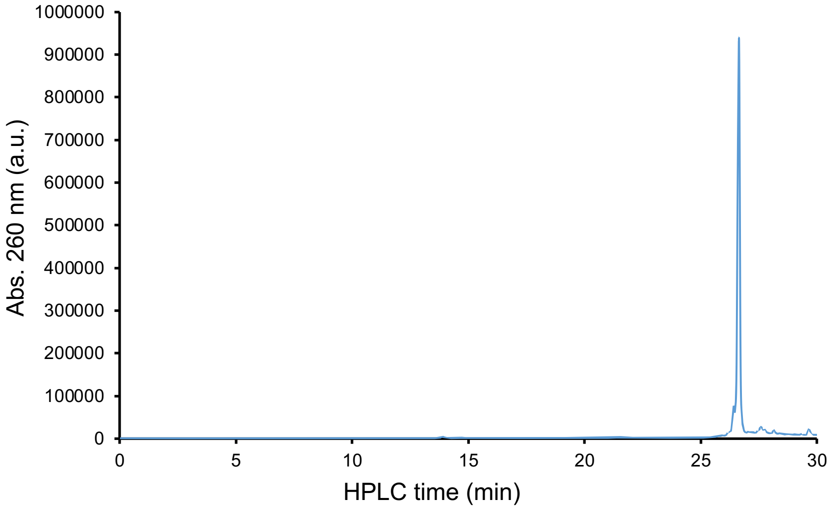


**A.**

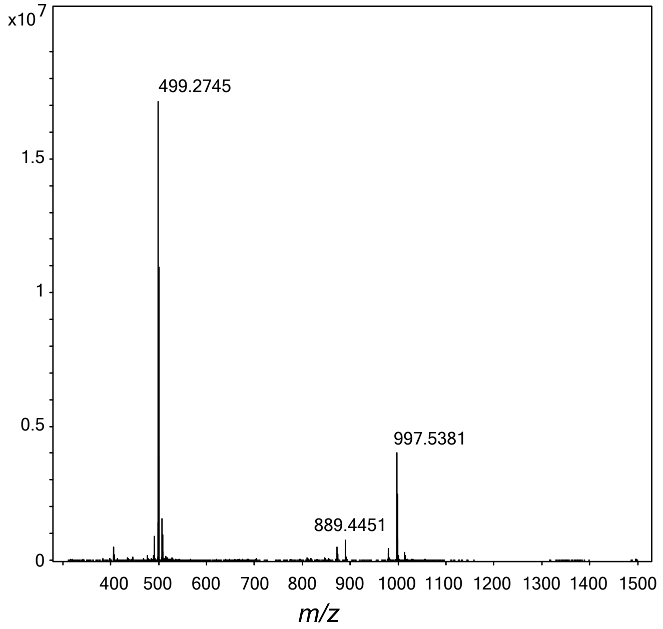


**B.**

**Figure S3.** Analytical HPLC (**A.**) and ESI-MS (**B.**) spectra of **TCO-CPP-RRWQW**.

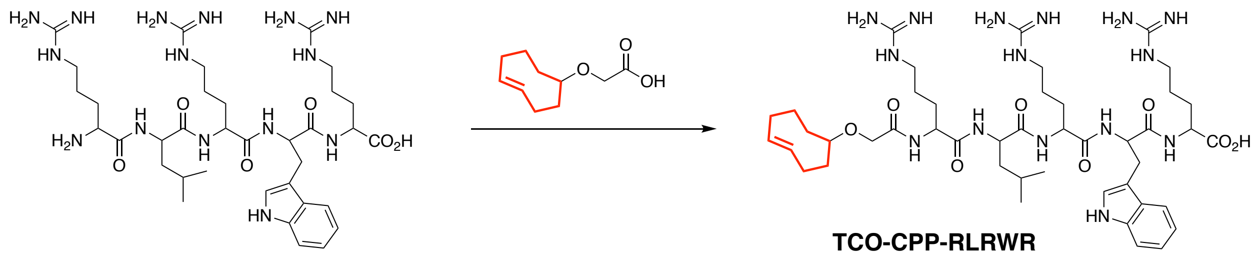


Prepared a suspension of HATU (14 mg) in DMF (200 μL). In parallel, prepared a solution of (*E*)-2-(Cyclooct-4-en-1-yloxy)acetic acid (3.3 mg) in NMP (100 μL). Combined the two solutions, added DIPEA (7μL) and 2,6-lutidine (7μL) and stirred for 10 min at rt. Added a solution of **RLRWR** (20 mg, 25 μmol) in DMF (100 μL) and stirred at rt for 2 h. **TCO-CPP-RLRWR** was purified by preparative HPLC and analyzed by analytical HPLC and ESI-MS, shown below. HRMS (ESI) cal'd for $C_{45}H_{74}N_{15}O_8$ $[M+1]^+$ 952.5839; observed 952.5846

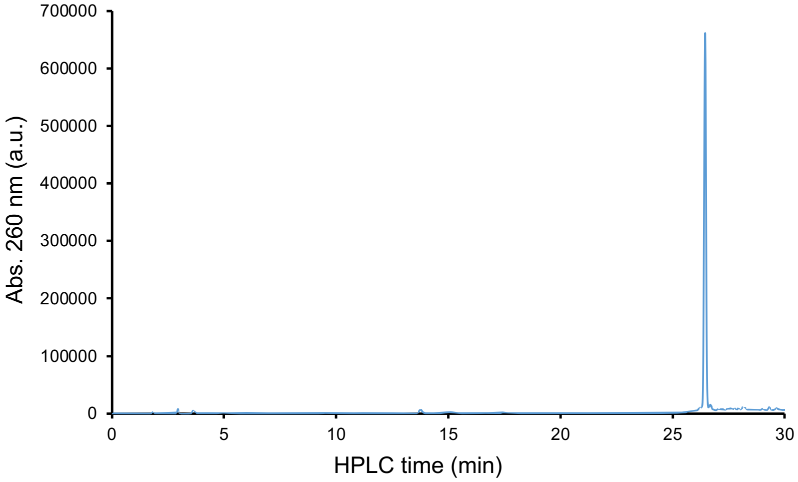


**A.**

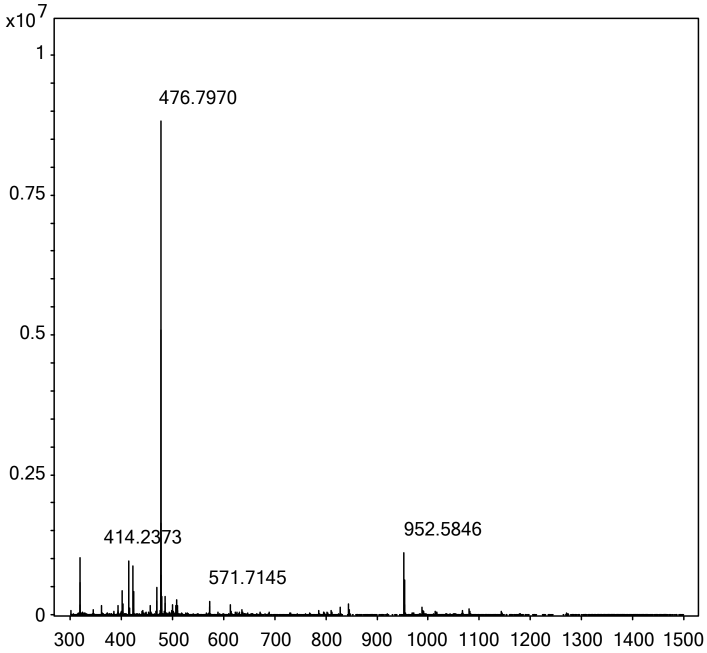


**B.**

**Figure S4.** Analytical HPLC (**A.**) and ESI-MS (**B.**) spectra of **TCO-CPP-RLRWR**.

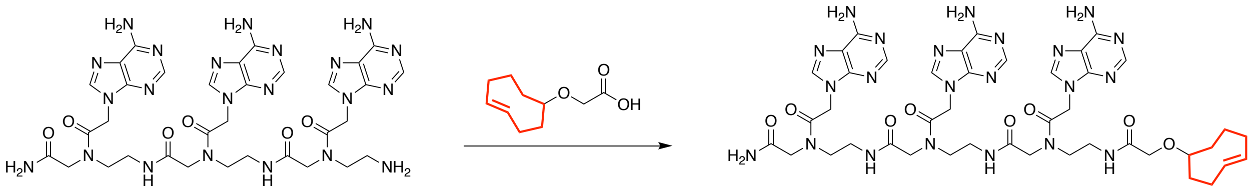


Prepared a suspension of HATU (14 mg) in DMF (200 μL). In parallel, prepared a solution of (*E*)-2-(Cyclooct-4-en-1-yloxy)acetic acid (3.3 mg) in NMP (100 μL). Combined the two solutions, added DIPEA (7μL) and 2,6-lutidine (7μL) and stirred for 10 min at rt. Added a solution of **AAA-PNA** (20 mg, 24 μmol) in DMF (100 μL) and stirred at rt for 2 h. **AAA-PNA-TCO** was purified by preparative HPLC and analyzed by analytical HPLC and ESI-MS, shown below.
HRMS (ESI) cal'd for $C_{43}H_{57}N_{22}O_8$ $[M+1]^+$ 1010.0745; observed 1012.0507

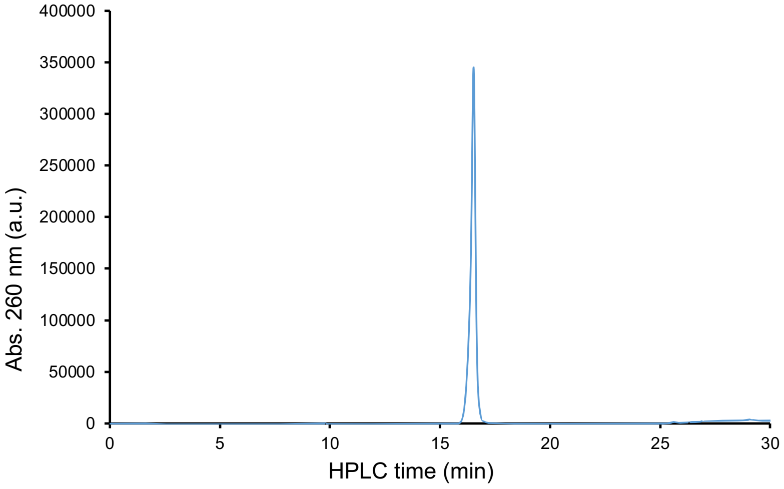


**A.**

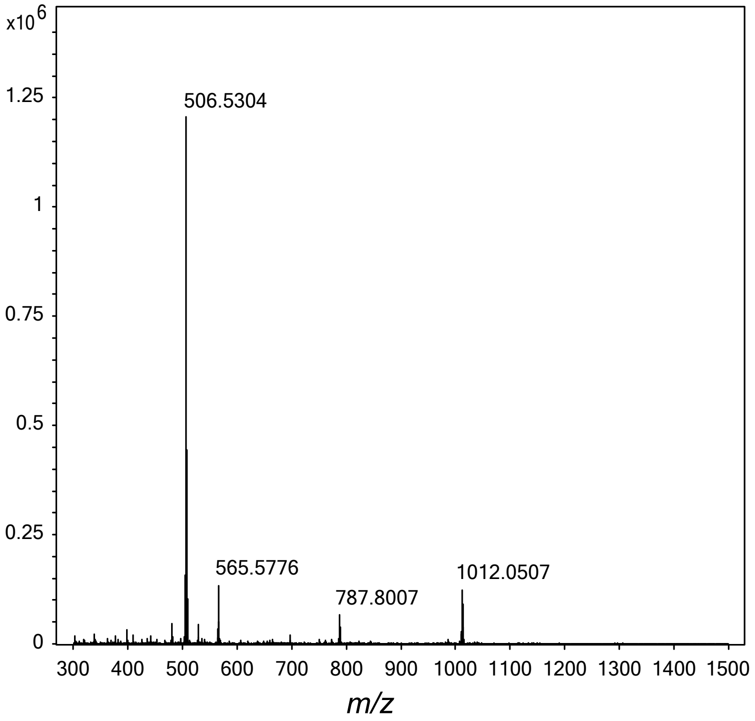


**B.**

**Figure S5.** Analytical HPLC (**A.**) and ESI-MS (**B.**) spectra of **AAA-PNA-TCO**.

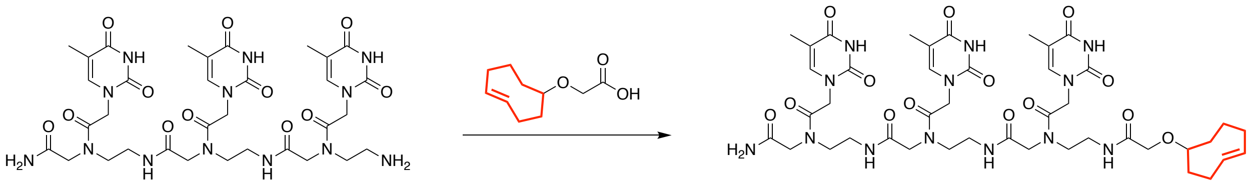


Prepared a suspension of HATU (14 mg) in DMF (200 μL). In parallel, prepared a solution of (*E*)-2-(Cyclooct-4-en-1-yloxy)acetic acid (3.3 mg) in NMP (100 μL). Combined the two solutions, added DIPEA (7μL) and 2,6-lutidine (7μL) and stirred for 10 min at rt. Added a solution of **TTT-PNA** (20 mg, 25 μmol) in DMF (100 μL) and stirred at rt for 2 h. **TTT-PNA-TCO** was purified by preparative HPLC and analyzed by analytical HPLC and ESI-MS, shown below.
HRMS (ESI) cal'd for $C_{43}H_{60}N_{13}O_{14}$ $[M+1]^+$ 982.4377; observed 982.4356

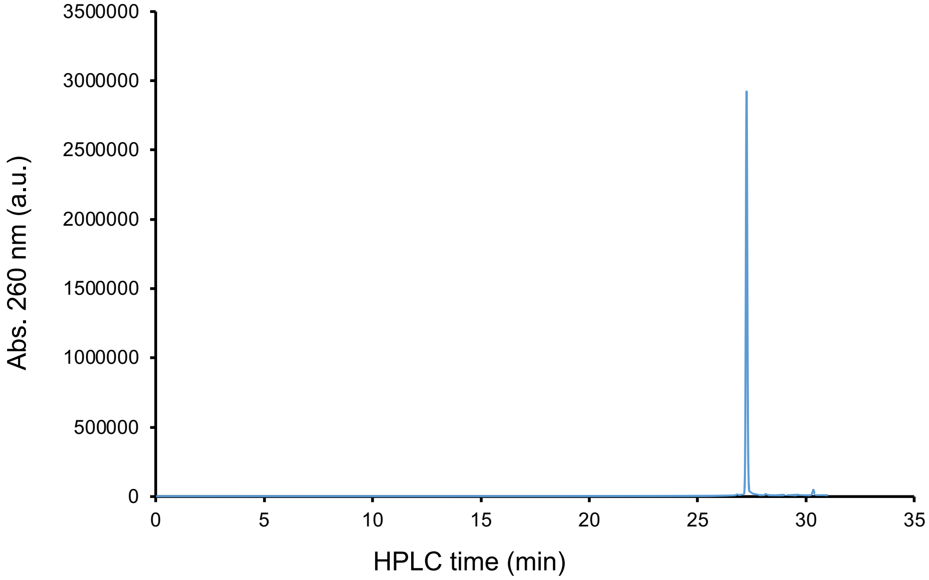


**A.**

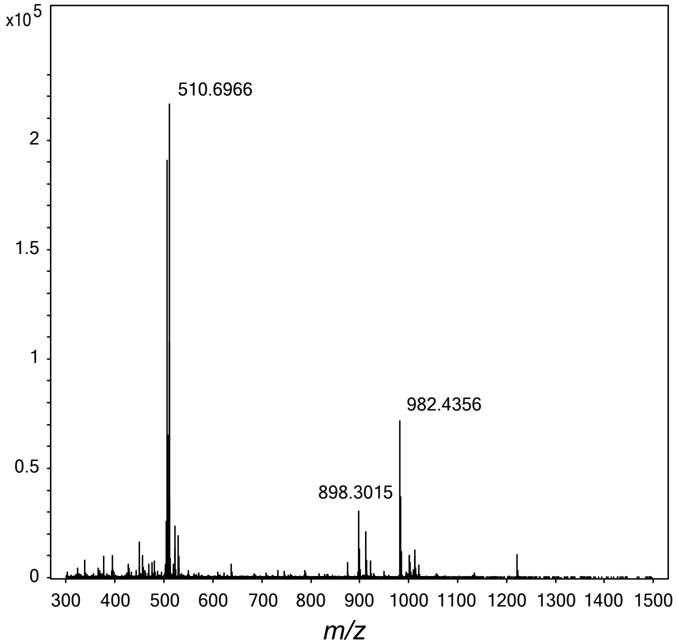


**B.**

**Figure S6.** Analytical HPLC (**A.**) and ESI-MS (**B.**) spectra of **TTT-PNA-TCO**.

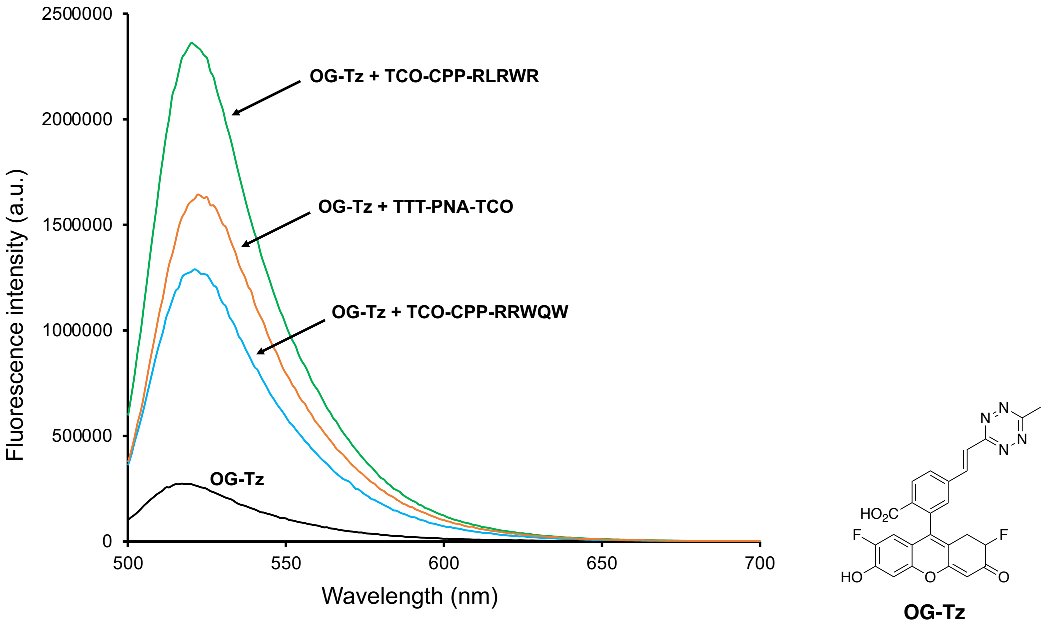


**Figure S7.** Fluorescence spectra of **OG-Tz** ($\lambda_{ex}$ = 490 nm) by itself and conjugated to different TCO-modified CRISPR suppressors.

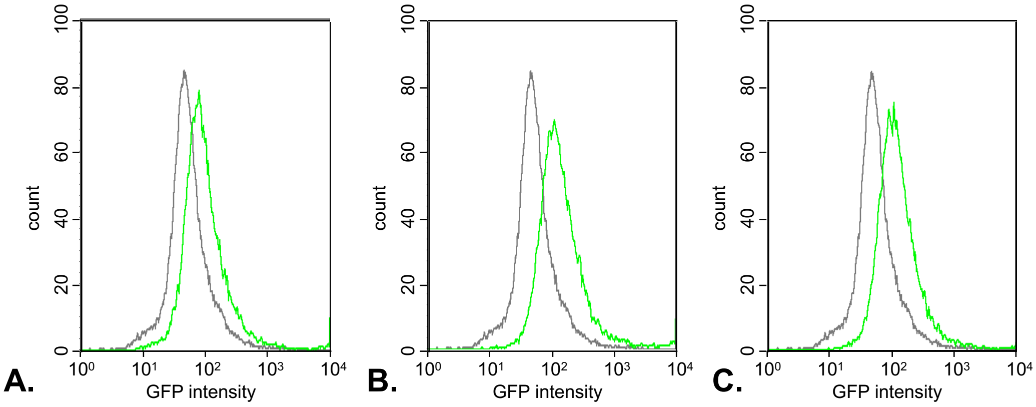


**Figure S8.** Flow cytometry of HEK293 cells treated with **OG-Tz** alone (black) and with **OG-Tz** and TCO-modified CRISPR suppressors (green). (A) **TTT-PNA-TCO**; (B) **TCO-CPP-RRWQW**; (C) **TCO-CPP-RLRWR**.

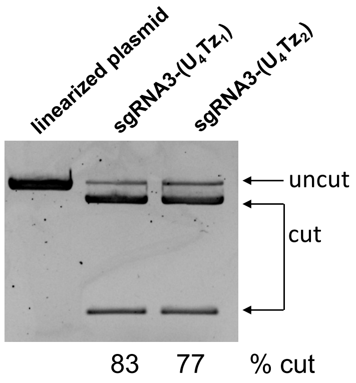


**Figure S9.** Analysis of CRISPR-Cas9 experiments using agarose gel electrophoresis. (**A**) *Lane 1*: linearized eGFP-N1 plasmid; *Lane 2*: linearized eGFP-N1 plasmid and **sgRNA3-($U_4Tz_1$)**; *Lane 3*: linearized eGFP-N1 plasmid and **sgRNA3-($U_4Tz_2$)**.
